# Combining crystallographic fragment screening, algorithmic merging and digitized chemical synthesis efficiently yields scaffolds that target diverse NCS-1 protein-protein interactions implicated in neurological disorders

**DOI:** 10.1101/2025.06.18.660085

**Authors:** Daniel Muñoz-Reyes, Kate K. Fieseler, Max Winokan, Mathew Golding, Eda Capkin, Matteo Ferla, Sara Pérez-Suárez, Charlie W. E. Tomlinson, Peter G. Marples, Celia Miró-Rodríguez, Alicia Mansilla, Daren Fearon, Warren Thompson, Frank von Delft, María José Sánchez-Barrena

## Abstract

Efficient drug discovery relies on workflows that integrate structural insights with rapid and cost-effective exploration of chemical space. Here, we present a data-driven fragment-based lead discovery approach to target Neuronal Calcium Sensor 1 (NCS-1) protein-protein interactions (PPIs). This study represents a complete implementation of a single high-value design-make-test-analyze cycle that directly yields compounds with micromolar affinity with the potential to modulate NCS-1 interactions with key targets, including the G-protein chaperone Ric-8A and the dopamine D_2_ and cannabinoid CB_1_ receptors. X-ray crystallographic fragment screening (CFS) revealed diverse interaction patterns within the NCS-1 hydrophobic crevice. Algorithmically guided fragment evolution and automated synthesis enabled the rapid generation of over 250 derivatives, with biophysical validation using LC-MS and Grating-coupled interferometry. Structural analyses highlighted key pharmacophores, with selected compounds exhibiting favorable drug-like properties and potential blood-brain barrier penetration, making them promising candidates for neurodegenerative and neurodevelopmental disorders. Our results demonstrate the feasibility of accelerated hit-to-lead development at synchrotrons, demonstrating a robust, scalable platform for PPI-targeting drug discovery. The generated chemically diverse scaffolds provide a strong foundation for future therapeutic optimization.

## INTRODUCTION

The Neuronal Calcium Sensor 1 (NCS-1) is a calcium signaling protein with multiple functions in the central nervous system and in other tissues^1^. NCS-1 interacts with several G-protein-coupled receptors (GPCRs) (adenosine A_2A_, cannabinoid CB_1_ and dopamine D_2_ receptor) and proteins implicated in regulating G-signaling, among others, the molecular chaperone and guanine exchange factor (GEF) Ric-8A^2^ or the kinase GRK2^3^. In addition, it also interacts with calcium channels such as InsP_3_ receptor, which regulate cytosolic calcium concentrations^4^.

NCS-1 interacts with its binding partners through a large solvent-exposed hydrophobic crevice, a structural feature shared across the Neuronal Calcium Sensor (NCS) protein family. The 16 protein members display up to 60% sequence identity, adopt similar overall topologies, and conserve key hydrophobic residues within the binding pocket that are critical for partner recognition. Structural analyses of several NCS-protein complexes indicate that binding specificity is dependent on the shape and dimensions of the cavity. The cavity is influenced by: (i) the Ca^2+^ content of the protein, (ii) the presence of hydrophilic residues at the border of the crevice, which serve as interaction hotspots; and (iii) the C-terminal and dynamic helix H10, which can insert into the cavity. Helix H10 contains an unstructured C-terminal tail that allows this region to adopt multiple conformations, thereby contributing to the shape of the crevice^5–7^.

NCS-1 has been implicated in disease pathogenesis^8^. In the past few years, it has been shown that the NCS-1/Ric-8A interface is a druggable target and the inhibition or enhancement of this protein-protein interaction has therapeutic relevance in neurodevelopmental disorders or neurodegeneration^5,6,9,10^. Given the large size of the protein-protein interaction (PPI) interface, and the multiple binding partners of the calcium sensor, it would be of value to fully explore the chemical space of this PPI and design compounds targeting different regions of the crevice to yield selective PPI regulators that allow the modulation of specific targets while avoiding a global effect on the NCS-1 interactome (Figure 1).

**Figure 1:**
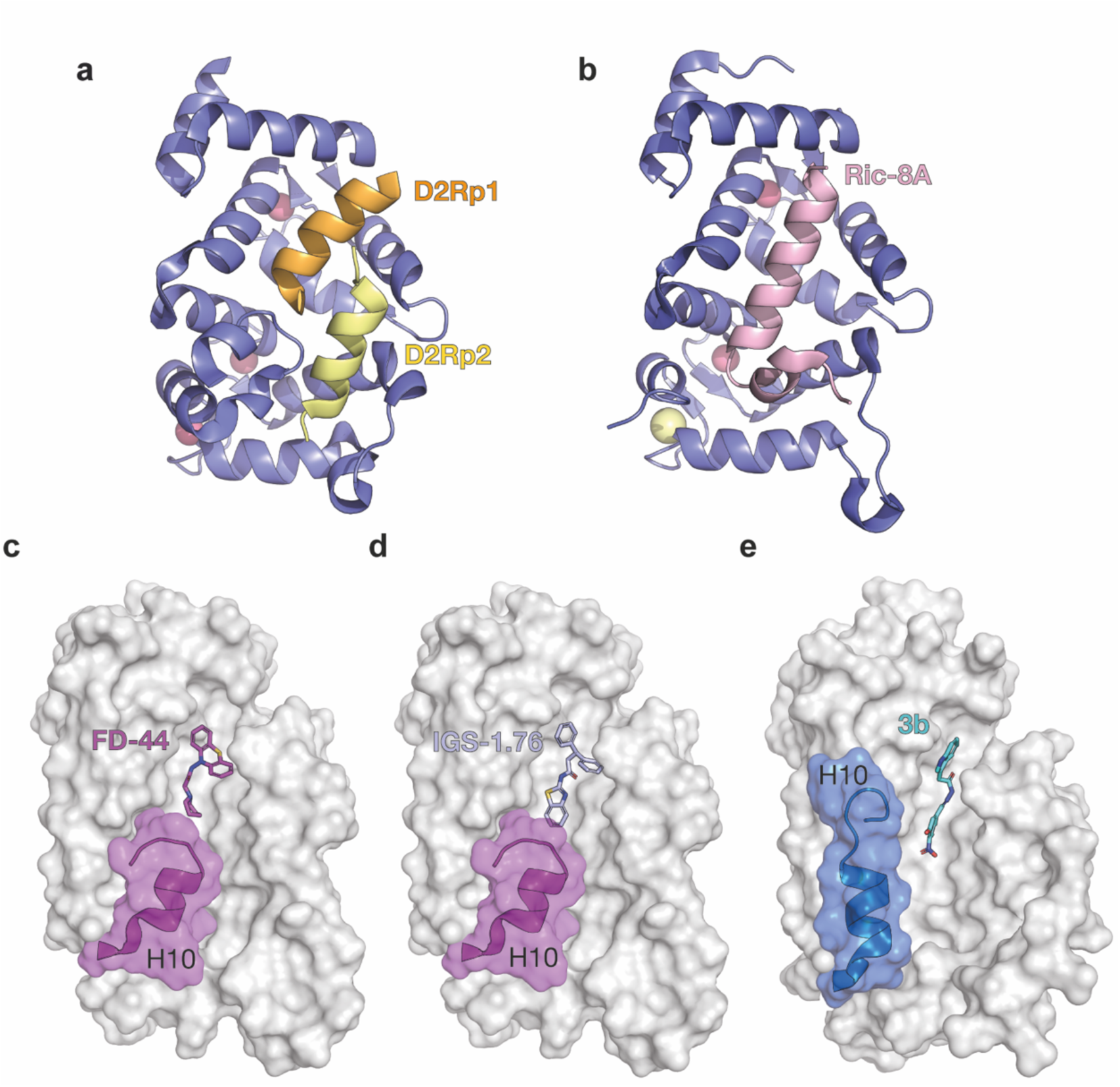
The structure of NCS-1 bound to target proteins and ligands complexes. (**a**) and (**b**) Ribbon representation of NCS-1 protein structures bound to a dopamine D_2_ receptor C-terminal peptide (PDB: 5AER,^11^) and the molecular chaperone and GEF Ric-8A (PDB: 8AHY,^7^), respectively. (**c**), (**d**) and (**e**) Structures of NCS-1 in complex with regulatory ligands, the NCS-1/Ric-8A PPI inhibitors FD-44 (PDB: 5AAN,^5^) and IGS-1.76 (PDB: 6EPA,^6^) and the PPI stabilizer 3b (PDB: 6QI4,^9^), respectively.

Over the past decade, the development of high-throughput crystallographic fragment screening facilities at synchrotrons has emerged as a powerful methodology for mapping protein binding sites and exploring the chemical space of these targets. This approach enables the identification of novel hits with significant biomedical and therapeutic potential. The XChem facility at Diamond Light Source has enabled screening of over 1,000 fragments per week, thanks to advances in crystal identification, automated crystal soaking, crystal mounting, automated data collection and crystallographic data processing^12,13^. This strategy requires a robust crystal system with highly reproducible crystallization, tolerance to the organic solvent used to dissolve the fragments, and ideally, diffraction at a resolution of better than 2.5 Å^12^. The true bottleneck in developing lead-like compounds from crystallographic fragment screens lies in the efficient exploitation and exploration of the structural information contained in protein-fragment complexes to guide structure-based drug design. Few existing procedures are both efficient and fully automated^14,15^. At XChem, algorithmic approaches have been developed to complete a comprehensive pipeline for managing the various stages of the DMTA (Design-Make-Test-Analyze) cycle, with specific focus on the Design, Make and Test components. Reaction purification is currently a complete blocker to high-throughput synthesis to biology readout; exploiting ligand-protein affinity to selectively bind to the desired compound in a crude reaction matrix is a viable approach to eliminating the purification bottleneck^16–18^. The XChem fragment progression workflow has been implemented to rapidly (<6 months) and at relative low-cost (€8k for building blocks and €2,6k for pure compounds) explore the chemical space of the NCS-1 PPI interface and develop chemically diverse molecules that allow the design of selective compounds against this pharmacologically interesting target (Figure 2).

**Figure 2:**
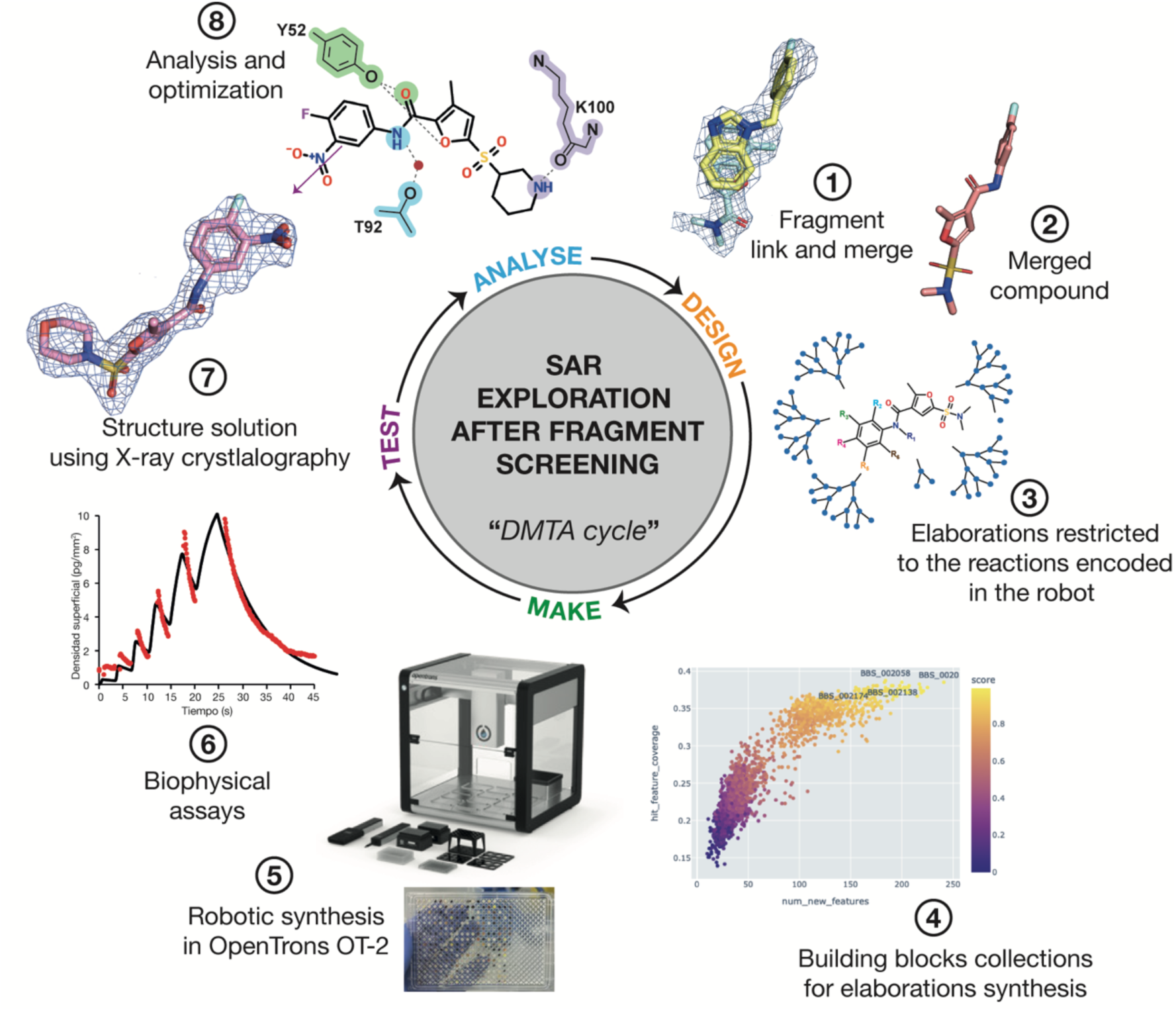
Implementation of the DMTA cycle in NCS-1. The DMTA cycle consists of four distinct phases: Design (orange), make (green), test (purple) and analyze (blue). The different steps (1-8) implemented in this study for the search of new modulators of the interaction between NCS-1 and its target proteins are shown.

## RESULTS

### Crystallographic fragment screening productively samples all subpockets

A previously reported full-length hNCS-1 crystal form^19^ diffracts beyond 2Å and tolerates DMSO making it suitable for large-scale fragment screening. Furthermore, its soakability was demonstrated in the structure of NCS-1 bound to theNCS-1/Ric-8A PPI stabilizer 3b (PDB: 6QI4,^9^). In the crystal, the NCS-1 hydrophobic crevice is exposed to the solvent, and the C-terminal helix H10 is positioned laterally, allowing compound entry into the cavity where the NCS-1 target proteins bind (Figure 1e).

The fragment libraries were selected for their chemical diversity and the presence of chemotypes compatible with the hydrophobic crevice of NCS-1. The York 3D library was especially attractive as it contains substituted aliphatic heterocycles, moieties that are contained in previous regulators of the NCS-1/Ric-8A PPI (Figure 1c to e)^5,6,9^.

A total of 1,258 crystals were mounted from 1,600 crystal drops and automated data collection carried out on the I04-1 beamline (Materials and Methods). Of the 937 datasets collected, only 540 were of sufficient quality and resolution to solve the corresponding protein-fragment structures (Figure 3). Hit identification with PanDDA has limited success for NCS-1, as several structural elements, including the unstructured N-terminal region and the dynamic C-terminal H10 helix, exhibited conformational changes upon soaking and fragment binding. Thus, hit identification was more challenging, and manual building of these dynamic elements was required to improve electron density maps and allow hit detection. Additional electron density was observed in 135 datasets, from which fragments could be modeled in 49, resulting in a total of 82 fragments (Figure 3a). The resolution of these 49 datasets ranged from 1.5 to 2.3 Å, with the majority falling between 1.7 and 1.9 Å (Figure 3d).

**Figure 3:**
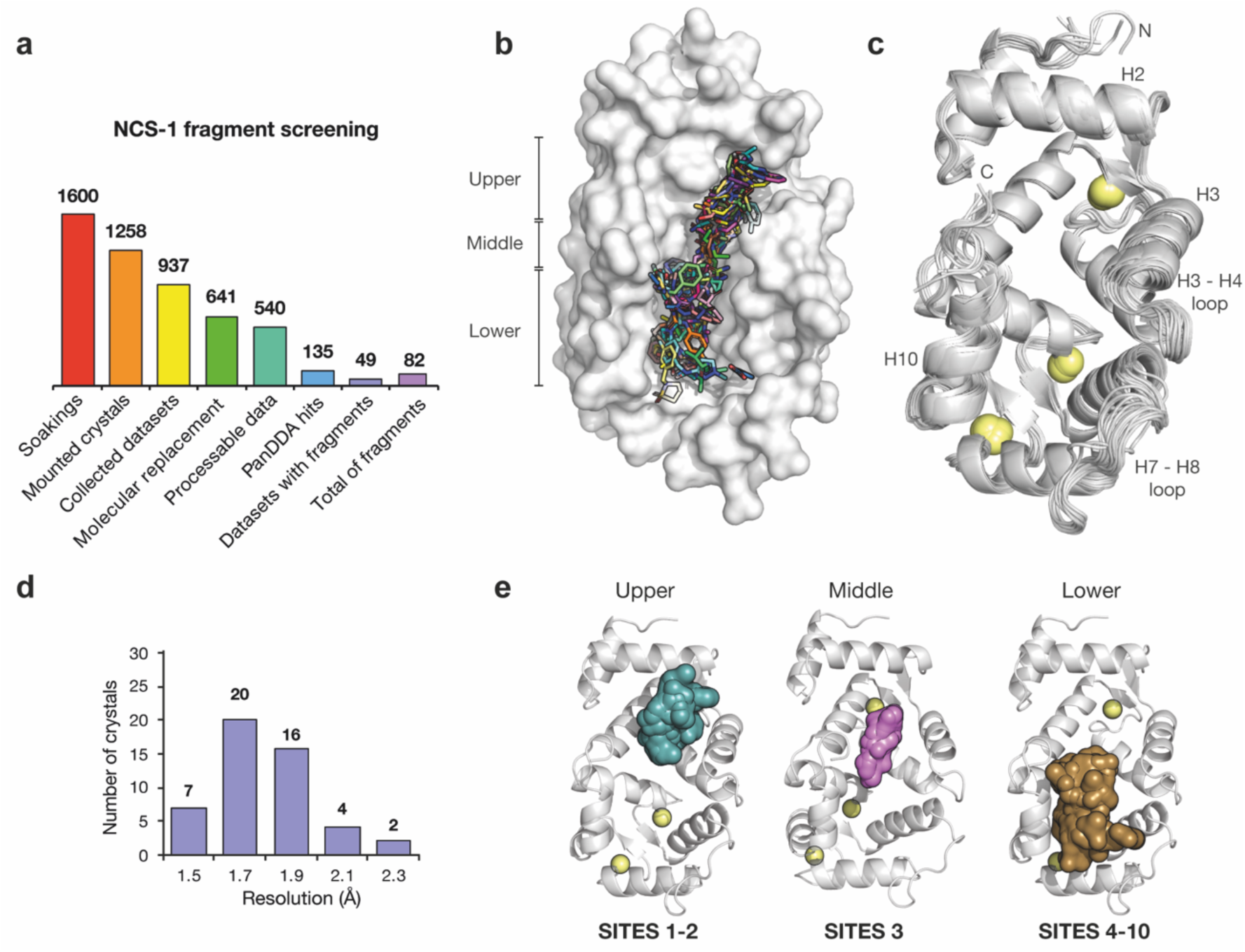
Fragment screening by X-ray crystallography identifies 83 fragments bound to NCS-1. (**a**) Histogram of the NCS-1’s fragment screening process. (**b**) Representation of the molecular surface of NCS-1 and the superposition of identified fragments (shown as sticks). (**c**) Superposition of NCS-1 fragment-bound structures. Ca²⁺ ions shown as yellow spheres. Variable regions are indicated. (**d**) Histogram displaying the maximum resolution of crystals where fragments were identified. (**e**) Distribution of fragments along the NCS-1 crevice. Fragments occupying the different regions of the crevice (upper, middle and lower part) have been represented as molecular surface in green, pink and brown, respectively. Fragment sites belonging to each region are indicated.

### The analysis of fragment distribution reveals the chemical space of the hydrophobic crevice of NCS-1

An overlay of all fragment hits shows that the entire hydrophobic crevice is occupied by ligands (Figure 3b). Some datasets revealed up to four distinct binding events along the NCS-1 crevice, while others showed only one. An overlay of the ligand-bound NCS-1 chains highlights regions with the greatest conformational variability upon ligand binding (Figure 3c). These include the N-terminal region of the protein, the dynamic helix H10 and its C-terminal unstructured tail, the Ca^2+^ binding loop of EF-hand 1 (EF-1), which is insensitive to Ca^2+^ binding, and the loops connecting EF-hands EF-2 and EF-3 (H3-H4 loop) and EF-3 and EF-4 (H7-H8 loop) (Figure 3c).

Fragments were observed to bind along three main regions of the hydrophobic crevice, the upper, middle, and lower region (Figure 3c and e). Based on the residues involved in fragment recognition, ten distinct binding sites were identified. Sites 1 and 2 are in the upper region of the cavity, Site 3 in the middle, while Sites 4 to 10 in the lower region (Figure 3e).

#### Fragments bound to the upper region of the NCS-1 cavity

A total of 20 unique fragments bind in the upper part of the cavity (Supplementary Figure 1). Site 1 contains eleven unique fragments that interact with helix H2 (Supplementary Figure 1a). All fragments establish contacts with residue W30 through π-π interactions or hydrophobic van der Waals contacts. Additionally, four fragments form hydrogen bonds with D37, while the remaining ten interact with D37 through van der Waals contacts (Supplementary Figure 1e). The fragment Z86416893 is of particular interest, as it is strongly anchored to helix H2 through two hydrogen bonds with D37. It also forms π-π interactions with W30 and further van der Waals contacts with nearby residues (Supplementary Figure 1b). These interactions make this fragment a promising candidate for further development. Site 2 contains nine unique fragments that interact with key residues on helices H3 (Y52) and H5 (F85 and T92) (Supplementary Figure 1c and d). These fragments can be divided into three subgroups based on their interactions: those that form only hydrophobic interactions with F85, those that form a direct hydrogen bond with Y52, and those that establish a direct hydrogen bond with T92 (Supplementary Figure 1f).

#### Fragments bound to the middle region of the NCS-1 cavity

A total of five fragments bind to the middle part of the NCS-1 cavity, which represents Site 3 (Supplementary Figure 2). In all cases, H-bonds are established with residues Y52 or T92 (Supplementary Figure 2c), which are considered hotspots in the modulation of the NCS-1/Ric-8A complex^5,6^. The fragment Z55290386 is considered highly promising as it is strongly stabilized with three hydrogen bonds, one direct bond with Y52 and two water-mediated bonds with T92 (Supplementary Figure 2b).

#### Fragments bound to the lower region of NCS-1 cavity

A significant proportion of the fragments, a total of 42 fragments, bind to the lower part of the NCS-1 cavity (Supplementary Figure 3 and Supplementary Figure 4). This is due to the orientation of helix H10, which generates a deeper cavity in this region. To date, no modulators have been reported to bind to this region. This lower part of the crevice has been subdivided into seven binding sites based on the relative positions of the fragments within this region (Supplementary Figure 3a to c and Supplementary Figure 4a to d). Fragments in Site 4 feature an aromatic group that forms hydrophobic contacts or π-π interactions with W103, located in the lower part of the cavity. A total of twenty-eight unique fragments spans the cavity (Supplementary Figure 3a and g). In addition, some fragments form hydrogen bonds with Y52 (Z31723394) or with various residues of helix H10. In contrast, the seven fragments in Site 5 are accommodated laterally; they do not establish H-bonds with helix H10 but make van der Waals contacts with W103 (Supplementary Figure 3b and h). The six fragments in Site 6 do not interact with W103 (Supplementary Figure 3c and i), but some form hydrogen bonds with Y108 or R148. The remaining fragments are positioned in Sites 7 to 10, in the deepest region of the NCS-1 cavity (Supplementary Figure 4). Fragments in Site 7 are placed between Y108 and R148 (Supplementary Figure 4a and e). Most of these fragments feature a protein-interaction aromatic group and contain amine groups or oxygen atoms to establish H-bonds with Y108 or R148, at the very end of the hydrophobic crevice. Fragments in Site 8 overlap with those in Site 7 (Supplementary Figure 4b and f); however, these five fragments displace the R148 side chain, opening the lower part of the cavity (Supplementary Figure 4b). Their interactions are similar to those observed in Site 7. The three fragments in Site 9 are located even deeper in the cavity (Supplementary Figure 4c and g), while Site 10 is represented by a single fragment that interacts with the dynamic loop connecting EF-3 and EF-4 (Supplementary Figure 4d and h). This fragment is a thiophene-2-carboxylate, which forms π-π interactions with Y129 and establishes hydrogen bonds, mediated by a water molecule, with residues N134 and V136 (Supplementary Figure 4d).

### Fragment merging guided by structural analysis of protein-protein interactions generates NCS-1 PPI modulators

Before modifying, linking, and merging ligands, it is important to consider the structure and dynamics of NCS-1, as well as protein-protein recognition mechanisms. In this crystal form, NCS-1 positions helix H10 outside the crevice, participating in crystal contacts, which leaves the hydrophobic pocket fully exposed. However, in solution, helix H10 is dynamic, moving in and out of the cavity, and its position is critical for NCS-1 recognition of its targets^7,11^. Therefore, the positioning of helix H10, and thus modulation of protein-protein interactions, could be regulated by fragment binding^5,6,9^. Additionally, the structural determinants of NCS-1 target recognition are well-known, such as in the case of D_2_R^11^ and Ric-8A^7^. Therefore, compounds could be designed to target key interaction sites or hotspots.

The 83 unambiguously modelled fragments were used for a fragment progression strategy based on merging and linking fragment-hit combinations using Fragmenstein^20,21^, along with the placement of commercially available analogs sourced from multiple vendors (Materials and Methods). The high number of fragment hits and their spatial proximity enabled the generation of over 25,000 merged compounds. Given the potential therapeutic application of the resulting compounds in central nervous system disorders, a molecular weight cut-off of 500 Da was applied to the merged compound designs. To further prioritize suitable candidates, the compounds were filtered, ranked and classified using the following criteria: (i) the root mean square deviation (RMSD) between each merged compounds and its progenitor fragments; (ii) the Tanimoto coefficient, to evaluate chemical similarity between merged compounds and fragments; and (iii) the change in theoretical binding free energy (ΔG), of binding calculated using Fragmenstein. Next, we analyzed the overlap of the protein-ligand interactions with the protein-protein interactions found in the complexes of NCS-1 with its targets, Ric-8A, the dopamine D_2_ and cannabinoid CB_1_ receptors (hereinafter D_2_R and CB_1_R, respectively). For Ric-8A and the dopamine D_2_ receptor, the crystallographic structures of the protein-protein complexes were available^7,11^, while in the case of the cannabinoid CB_1_ receptor, an AlphaFold model was generated^22^ using available information on the region that the receptor uses to recognize NCS-1 (Figure 1) (Materials and Methods)^23^. Previous experimental structural data on modulators of the NCS-1/Ric-8A complex were also considered (Figure 1). Following these criteria in an iterative manner, the compounds were ultimately shortlisted through visual inspection. Finally, compounds were classified according to their potential to regulate each of the three PPIs by analyzing the overlap between protein-ligand and protein-target interaction sites^5,6,9^. Based on this analysis, the fragments that bound to the upper region of the NCS-1 crevice (Site 1 and 2) were classified as potential regulators of the NCS-1/D_2_R interaction. Modulators of the NCS-1/Ric-8A interaction were hypothesized to bind to the upper and middle parts of the crevice (Site 1 to 5). For the regulation of the NCS-1/ CB_1_R interaction, the fragments placed at Site 5 to 10 were selected, with the aim of targeting a different region and achieving selectivity. Furthermore, fragments that bound to the loop connecting EF-hands EF-3 and EF-4 or in the surrounding area were selected as this region is relevant for target recognition, is not conserved in the family of Neuronal Calcium Sensors and has not been explored before.

This structural analysis not only enabled the selection of sites for fragment pairing but also established a priority list of 66 compounds. Of these, 17 could be potential D_2_R modulators, 16 Ric-8A inhibitors, 16 Ric-8A stabilizers, and 13 CB_1_R modulators. Finally, there were 4 ligands which bound to the dynamic loop connecting EF-hands EF-3 and EF-4, that were selected.

17 out of the 66 compounds were commercially available and were purchased directly (Supplementary Table 1). They were chemical diverse and based in their predicted position within the binding pocket, they could potentially modulate different NCS-1 targets. Five, eight and four potential modulators of D_2_R, Ric-8A, and CB_1_R, respectively, were chosen (Supplementary Table 1 and Supplementary Figure 5).

### Algorithmic methods generate combinations compatible with robotic synthesis and SAR studies

The algorithmic method Syndirella (Synthetically Directed Elaborations) (https://github.com/kate-fie/syndirella) was employed to generate and select base compounds or “mergers” that could be used for SAR (Structure-Activity Relationship) studies and for designing elaborations that allow exploration of chemical space. Syndirella also ensures that the selected compounds are suitable for CAR (Chemist Assisted Robotics) robotic synthesis (Materials and Methods). Initially, the previously selected 66 base compounds were considered. Based on factors such as reaction selectivity, molecule growth vectors, and the cost of starting materials, 12 base compounds were ultimately selected and input into Syndirella. From these 12 mergers, Syndirella produced elaborated compounds designed to broadly explore the chemical and structural space of the NCS-1 hydrophobic crevice, while limiting the chemical synthesis to a single step. The proposed reactions included amidation, Schotten-Baumann sulfonamidation, Williamson synthesis, Suzuki reactions, and Chan-Lam coupling (Materials and Methods).

**Table 1:** Selected base compounds for elaboration and synthesis via CAR. The table shows the fragments that inspired the elaboration, predicted data from Fragmenstein and HIPPO, the synthesis summary, the products detected using MS and the number of crystal structures solved from crude extracts. (S) indicates the Site where the fragments of inspiration are found in the fragment screening.

|  | Base compound (Syndirella) |  |  |  |  |  |
| --- | --- | --- | --- | --- | --- | --- |
|  | D2R-5 | D2R-12 | CB1R-5 | Ric8inhib-1 | Ric8inhib-3 | Ric8enhan-3 |
| **Fragment inspiration** | Z86416893 (S1) + Z239127534 (S3) | Z53834613 (S1) + Z198195770 (S2) | Z126581390 4 (S4) + Z127248009 1(S6) | Z26333448 (S2) + Z126581390 4 (S3) | Z1265813904 (S3) + Z1198177230 (S4) | Z802824098 (S3) + Z1269220427 (S4) |
| **Synthesis reaction** | HATU amidation | HATU amidation | HATU amidation | HATU amidation | HATU amidation | Schotten-Baumann sulfonamidation with amine |
| **Fragmenstein** |  |  |  |  |  |  |
| $\Delta\Delta G$ (kcal/mol) | -11.0 | -9.5 | -11.7 | -9.2 | -8.2 | -11.7 |
| RMSD (Å) | 0.99 | 0.83 | 0.45 | 0.95 | 0.83 | 0.45 |
| Tanimoto | 0.19 | 0.22 | 0.46 | 0.25 | 0.28 | 0.46 |
| **Prediction of key interactions** |  |  |  |  |  |  |
| W30 | X | X |  |  |  |  |
| D37 | X | X |  |  |  |  |
| Y52 |  | X |  | X | X |  |
| T92 |  | X |  | X |  | X |
| K100 |  |  |  |  |  |  |
| W103 |  |  | X |  | X | X |
| Y108 |  |  | X |  |  | X |
| R148 |  |  | X |  |  |  |
| **Predicted Elaborations (HIPPO)** | 2 | 60 | 21 | 450 | 18 | 39 |
| **Synthesized elaborations (CAR)** | 0 | 27 | 14 | 383 | 14 | 16 |
| **Automated detection of products (MSCheck)** | - | 5 | 0 | 222 | 10 | 15 |
| **Synthesis success (%)** | - | 18.5 | 0 | 58.0 | 71.4 | 93.8 |
| **Crystal Structures** | - | 0 | 0 | 9 | 2 | 3 |

From the 12 proposed base compounds and over 5,000 successful elaborations, HIPPO (Hit Interaction Profiling for Procurement Optimization, https://github.com/mwinokan/HIPPO) was used to select those that would maximize the yield from the starting materials and produce the greatest number of elaborations within a fixed budget. These base compounds were named D2R-5, D2R-12, CB1R-5, Ric8inhib-1, Ric8inhib-3, and Ric8enhan-3; based on the PPI they were hypothesized to modulate (Figure 4, Supplementary Figure 6 and Supplementary Figure 7). The synthesis of all compounds was designed to proceed via a single synthetic step, primarily through an amidation reaction. An exception is Ric8enhan-3, which was synthesized using Schotten–Baumann sulfonamidation with amines (Table 1). Figure 4 illustrates the chemical space explored by the proposed base compounds and the elaborations designed, within a budget of €8,000. The selected collection of starting materials included the generation of 590 products from 280 starting materials. While the six base compounds cover 38.5% of the fragment screening interactions, the 590 products recapitulate 100% of the fragment interactions and introduce 241 new interactions (Figure 4).

**Figure 4:**
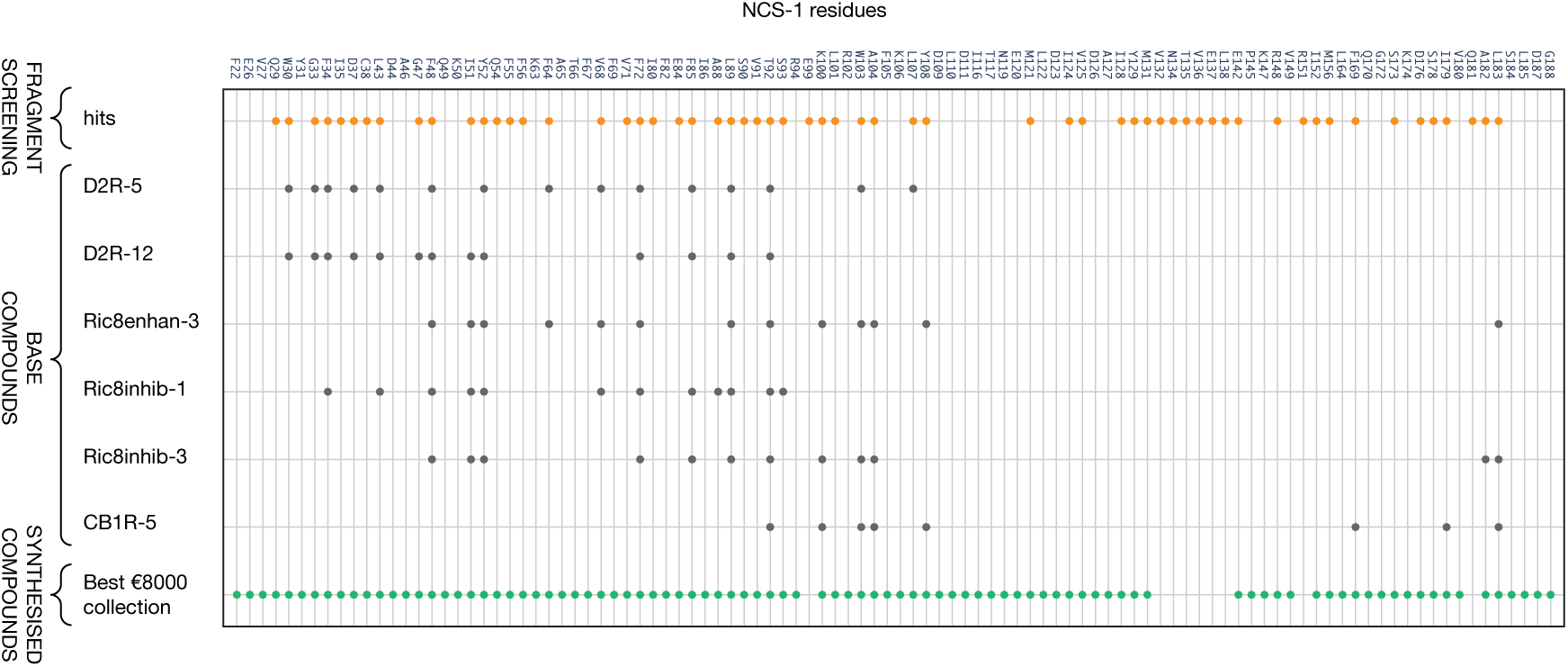
Exploration of the chemical space of NCS-1. Observed interactions in the NCS-1 fragment screening (hits; orange circles) with each NCS-1 residue. Predicted interactions for the six base compounds and the chosen collection of elaborations designed around the base compounds, suggested by Syndirella and adjusted to the €8000 reactant budget using HIPPO.

### New fragment-based compounds are efficiently generated via automated synthesis and validation

The automated synthesis of compounds was carried out in four different batches (Figure 5). Of the 590 reactions initially selected by HIPPO, 454 were successfully performed, representing 76.9% of the collection (Table 1).

Among the successfully performed reactions, elaborations based on Ric-8A-mod-1 were the most represented, accounting for 95.3% of the total elaborations. This prevalence is due to two main factors: first, the starting materials for Ric-8A-mod-1 elaborations were more affordable, and second, there was a large variety of available starting compounds, allowing for multiple combinations. Unfortunately, no elaborations based on D2R-mod-1 were synthesized, as the required starting compounds were unavailable at the time of purchase.

After synthesis, mass spectrometry was used to analyze the crude reaction mixtures; to facilitate the time-consuming task of analyzing hundreds of chromatograms. An automated LC-MS analysis tool called MSCheck^24^ was employed. In summary, MSCheck detected the presence of 15 sulfonamides and 237 amides, indicating that a total of 252 products had been synthesized. Therefore, the success rate of the sulfonamidation reactions was 93.8%, the success rate of the amidation reactions was 54.1%, and the overall success rate was 55.5% (Table 1 and Figure 5).

**Figure 5:**
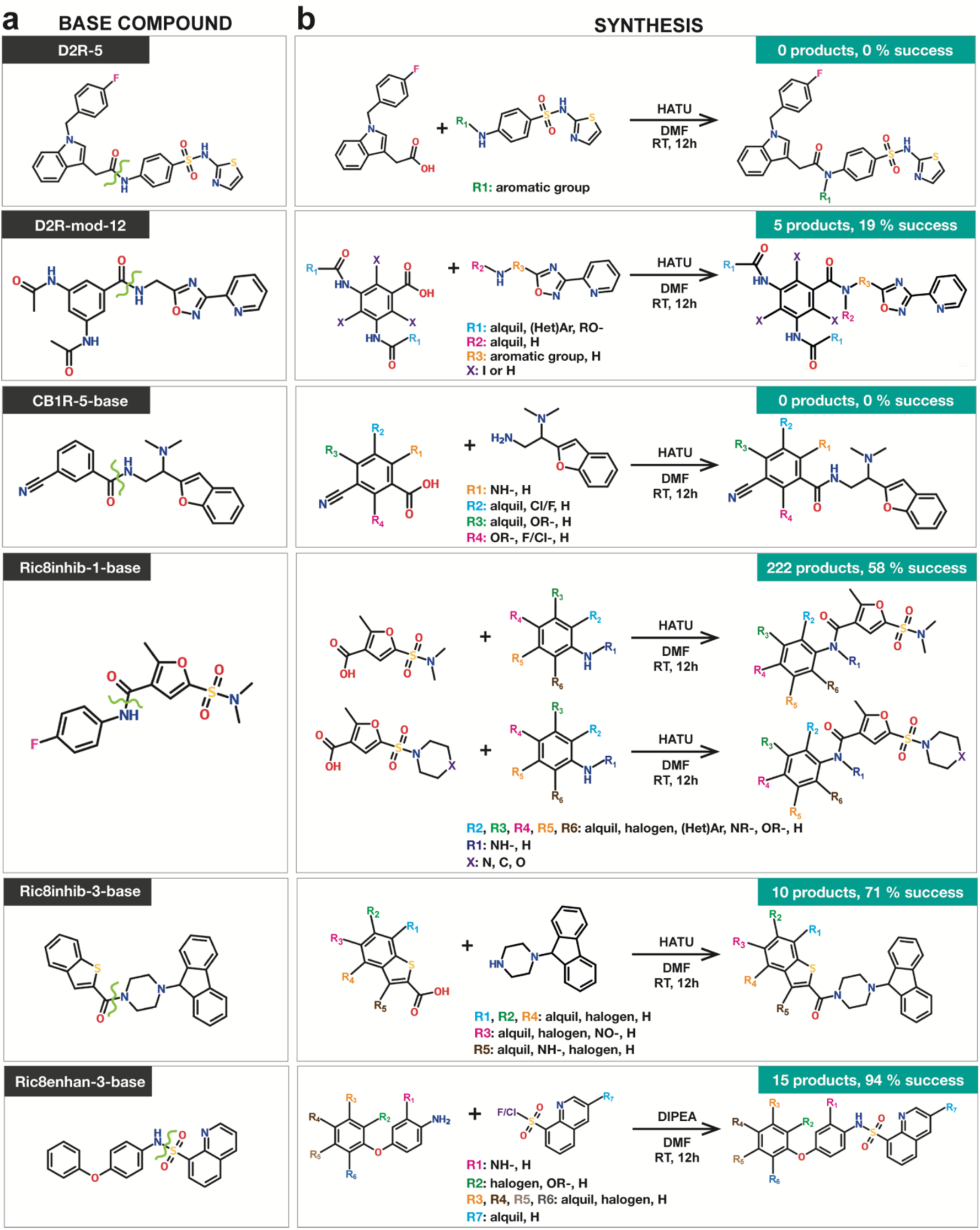
Fragment-based compounds synthesis via CAR. (**a**) 2D structure of the base compounds used for chemical elaboration. The green curve shows the bond created after merging the two starting molecules. (**b**) Synthesis and reaction conditions for each base compound. The starting materials are displayed along with the various functional groups (Rn, n=1-7). For each panel, the number of elaborated products and the success rate validated via LC-MS are indicated in the green box.

### Grating-coupled interferometry facilitates high-throughput characterization of compound affinities in solution

Grating-coupled interferometry (GCI) has been established at XChem as the orthogonal method for high-throughput determination of binding affinities. To detect the binding from impure and complex mixtures, a robust protocol was established first by studying the affinity of the purchased pure compounds, that served as a control. Six out of the seventeen pure compounds were detected as NCS-1 binders, representing 35% of the total compounds tested (Figure 6a and Supplementary Table 1). Under the experimental conditions, five of these compounds exhibited affinities for NCS-1 in the low micromolar range (E10, E12, E14, E16, and E17), while compound E6 demonstrated affinity in the tens of micromolar range (Figure 6a). Compounds E3 and E4 exhibited non-specific binding to the NTA surface and the remaining compounds showed no signal. In parallel, the affinity of NCS-1 for the compounds was also studied using tryptophan emission fluorescence to cross-validate the GCI results (Supplementary Figure 10). Agreement between the two techniques was observed, demonstrating that the immobilization of NCS-1 through a N-terminal 6xHis tag has no penalty on affinity measurement (Supplementary Table 2).

As previously mentioned, the primary goal of implementing this technique is to detect interactions of impure compounds from reaction mixtures. The sample concentration is needed to obtain affinity/kinetic information through the SPR or GCI strategy. Since the concentration of these mixtures is unknown, dissociation rate constants (k_d_) can be used for the evaluation of the binding interaction^25,26^. The off-rate screen is evaluated by fitting the dissociation phase for the ligand and sample interaction. The dissociation decay is independent of the sample concentration. The adjustment step was applied similarly to that used for the pure Enamine samples. Fitting parameters were adjusted based on the strength of the interaction between the sample and the target. Compound E17, with an affinity (K_D_) of 140 μM and a dissociation rate constant (k_off_) of 1.25 s^-1^, was used as a control (Figure 6a). In this study, the control compound E17 and other crude reaction mixtures have a fast dissociation profile. For this reason, the dissociation fit point was adjusted to 26.5 seconds. Among the 450 crude reaction mixtures tested, 15 samples indicated repeatable binding response and k_d_ values in the 1 and 2 s^-1^ range. Two of them (Ric8inhib-1-1111 and Ric8inhib-1-827) showed structure and k_d_ data (Figure 6b). By comparing dissociation kinetics, it is possible to detect compounds that can interact with NCS-1.

**Figure 6:**
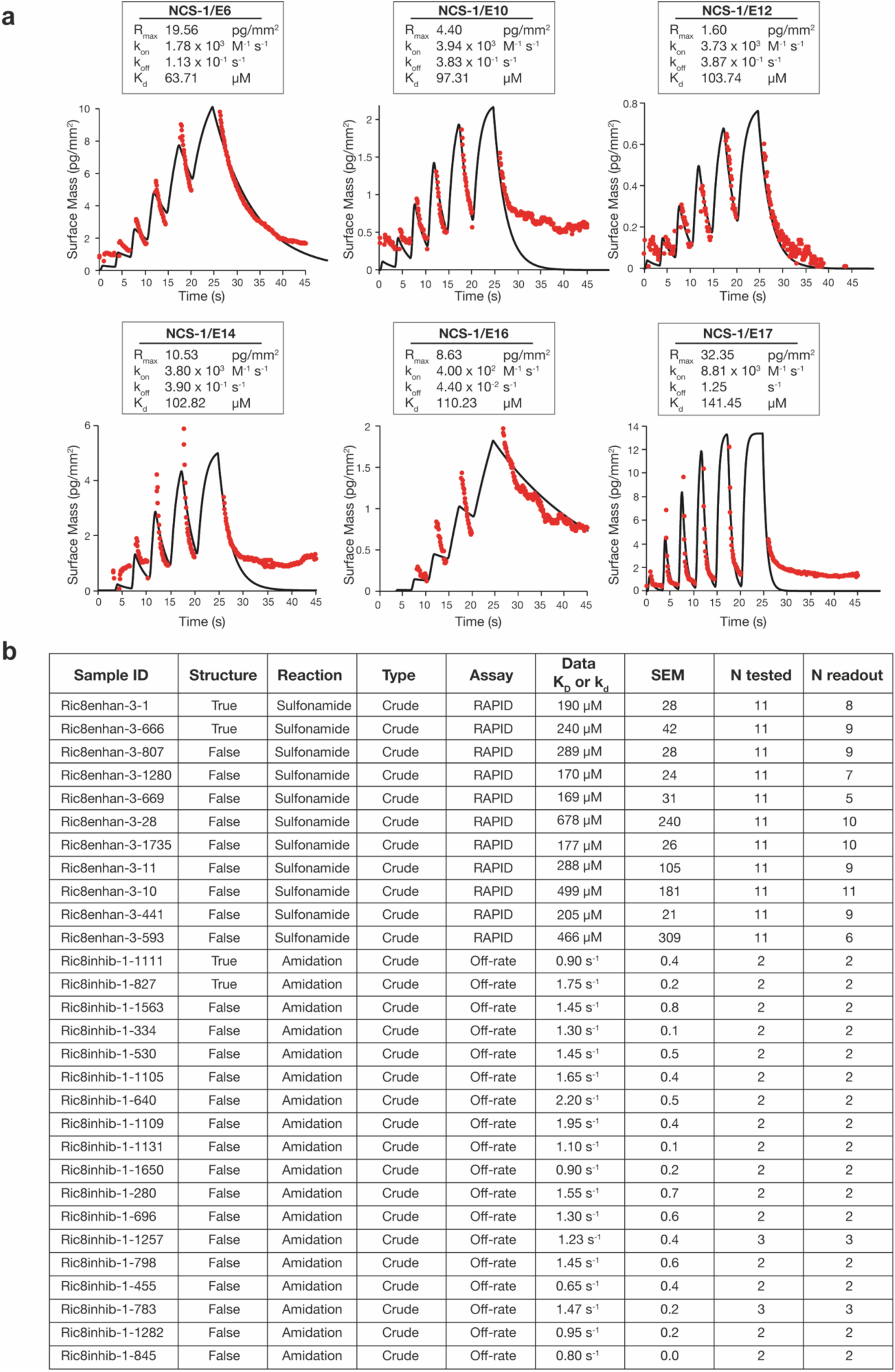
Study of the interactions between pure compounds and NCS-1 using grating-coupled interferometry (GCI). (**a**) Sensograms for pure compounds. Raw data are represented as red circles, with their respective fits in black. The binding kinetics were fitted to a 1:1 binding model. The tables summarize the kinetic parameters for each ligand: R_max_ (surface density of captured protein), k_on_ (association rate constant), k_off_ (dissociation rate constant), and K_d_ (dissociation constant). (**b)** Measurement of compounds from crude reaction mixtures. For each ligand, sample ID, crystal structure, reaction, assay, Kd or kd data and SEM (standard error of the mean) are indicated.

Product conversions were estimated from the LC-MS chromatograms and used by the RAPID binding assay to assess their kinetic parameters (11 measurements in total; Figure 6b). Among these, Ric8enhan-3-1 and Ric8enhan-3-666 showed both binding in crystals and measurable affinity. MSCheck data were used to estimate product concentrations, enabling more reliable affinity values. These preliminary data will help prioritize compounds for purification and further evaluation of their ability to modulate NCS-1 PPIs. However, it is important to note that crude reaction mixtures may contain residual reagents that compete with the products for NCS-1 binding, and thus the reported values should be considered indicative, not definitive.

### Crystallographic screening of lead-like compounds from pure and crude reaction mixtures and structural analysis

Previous studies have demonstrated that crude reaction mixtures, containing both starting materials and products, can be directly soaked into crystal drops, enabling the resolution of crystal structures of the synthetic products bound to the target protein without prior purification^16^. Initial experiments were performed similarly to the fragment soaks (2h, 10% DMSO). However, only one structure was solved. Therefore, longer incubation times, from 8 to 18h, were carried out. This resulted in 13 structures from of the 252 crude reaction mixtures. In addition, pure compounds were soaked and two structures of NCS-1 in complex with molecules E1 and E14 were solved.

In total, the soaking campaign led to the elucidation of 16 structures: 14 from CAR synthesis and 2 from the pure compounds (Figure 7 and Supplementary Figure 8). Both 2F_o_-F_c_ and F_o_-F_c_ electron density maps indicated the presence of ligands corresponding to the synthesized products via CAR occupying the NCS-1 hydrophobic crevice (Supplementary Figure 9). Except for compound Ric8-inhib-1-1111, where two ligands were bound to a NCS-1 molecule, all structures show 1:1 stoichiometry. In some cases, the quality of the density only allowed to model the ligand in one of the independent NCS-1 molecules. The corresponding poses observed in the two independent molecules were virtually identical except in compound E1, where the ligand binds to different cavities of the elongated NCS-1 hydrophobic crevice (Figure 7 and Supplementary Figure 8 and 9).

In the case of the structures obtained via CAR synthesis, the molecules are related to three base compounds, all of which were hypothetically designed to modulate the interaction between NCS-1 and Ric-8A (Supplementary Figure 7 and Table 1). The elaborations are located at the top and central region of the hydrophobic crevice of NCS-1 (Figure 7 and Supplementary Figure 8), distributed across the fragment Sites 1 to 5 (Supplementary Figure 1, Supplementary Figure 2 and Supplementary Figure 3). No structures were obtained from other elaborations hypothetically designed to modulate D_2_R and CB_1_R PPIs. However, it is worth noting that the number of molecules synthesized to potentially modulate the interaction with Ric-8A was much higher compared to those designed for D_2_R or CB_1_R (Table 1).

**Figure 7:**
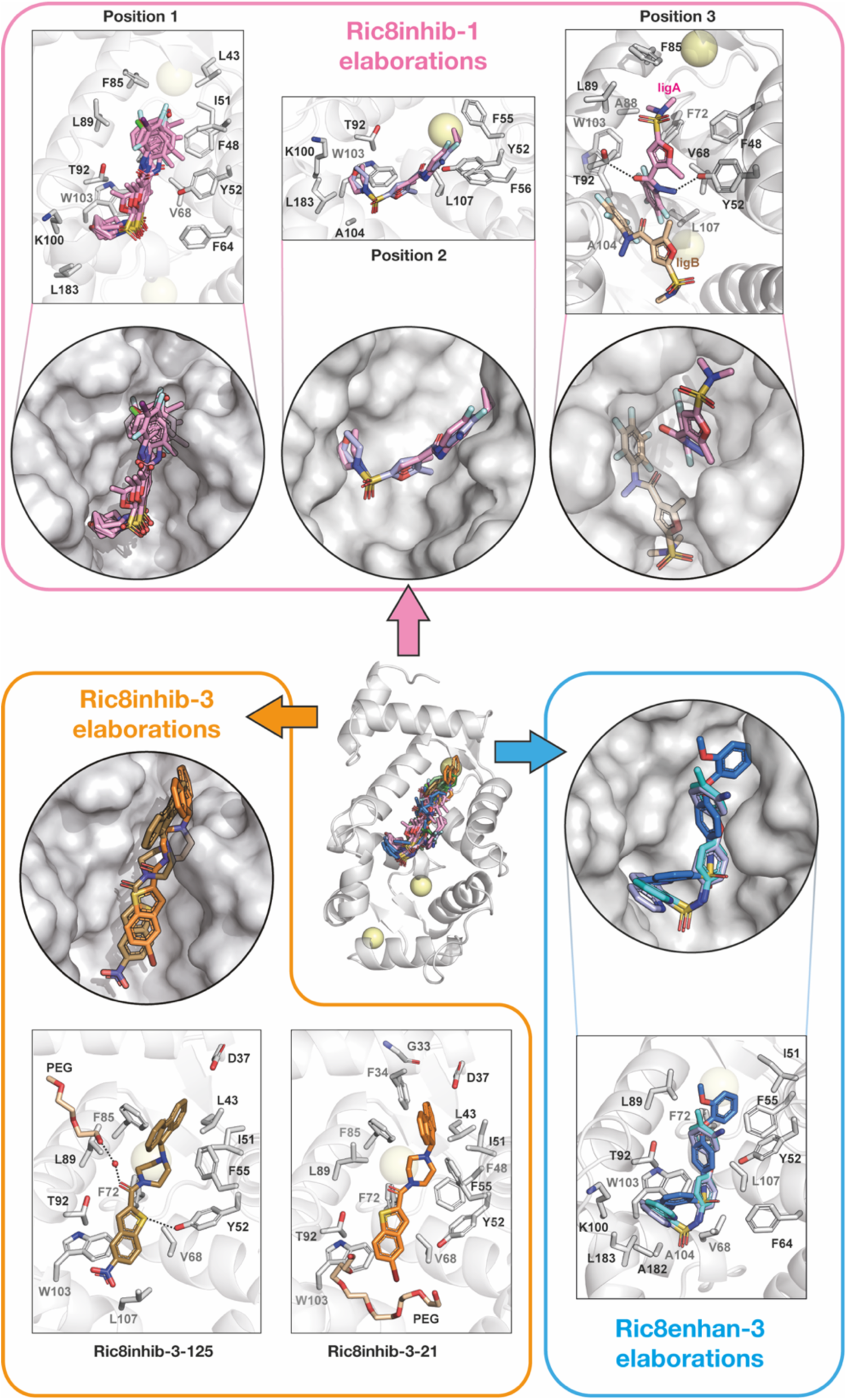
Screening of larger compounds via X-ray crystallography. In the center, structure of NCS-1 (ribbons) in complex with all the crystallized compounds (sticks), which have been superimposed. A magnification of the crystallized compounds Ric8inhib-1 (pink), Ric8inhib-3 (orange) and Ric8enhan-3 (blue) is displayed in each box. The NCS-1 cavity where ligands are located together with details on the interacting residues are also shown.

Nine structures were obtained from elaborations based on Ric8inhib-1 (Table 1, Figure 7 and Supplementary Figure 8). The products exhibit three different binding modes within the NCS-1 crevice. All the elaborations share a common scaffold based on 4-fluorophenyl-2-methyl-1-sulfonyl-3-furamide (Supplementary Figure 8b), differing chemically by the substituents on the 4-fluorophenyl group at one end of the ligand. Additionally, the sulfonyl group at the other end is substituted with either a dimethylamine or various non-aromatic nitrogen heterocycles (Supplementary Figure 8b). Most elaborations are in the intermediate part of the NCS-1 cavity (Position 1), where the amide group establishes key contacts with Y52 and T92 (Supplementary Figure 8b). These six ligands feature a heterocycle attached to the sulfonyl group at the lower part of the molecule, while no structures were resolved for elaborations with dimethylamine substitutions in this orientation, underscoring the importance of the heterocycle for NCS-1 interaction (Figure 7). At the top, they display an aromatic ring with various ortho, meta, and para substituents, forming hydrophobic contacts with neighboring residues in the cavity (Figure 7). The different substituents confer chemical diversity and allow the exploration of interactions with residues in the NCS-1 crevice. In addition, three elaborations were found to occupy other sites, termed Position 2 and 3 (Figure 7). Two ligands were located at Position 2, “floating” in the cavity and establishing predominantly hydrophobic contacts with aromatic residues in the base of the cavity (Figure 7). One ligand, Ric8inhib-1-1111, was identified in Position 3. This ligand features a dimethylamine group instead of the heterocycle present in the other compounds and a substituted amine on the furamide ring. Two molecules were found in the cavity, establishing contact between them. The presence of the amine group allows the molecule found in the upper region of the cavity to mediate important interactions in the central region of the crevice, orienting the ligand opposite to the other elaborations. The second molecule is found below, and p-stacking contacts are established through their furamide rings (Figure 7). Among all the compounds, Ric8inhib-1-1287 is of particular interest as it forms H-bonds with Y52 and T92, two key hotspots in the interaction with Ric-8A (Table 2). Furthermore, this ligand has a higher ΔG value than the other elaborations and forms numerous contacts with the protein. Conversely, the substitution of morpholine or piperidine groups with a pyrrolidine (elaboration 655) results in a lower ΔG value, indicating the importance of these groups for the interaction (Figure 7 and Table 2).

Two structures were solved with elaborations derived from Ric8inhib-3 which are bound in the upper half of the NCS-1 crevice (Figure 7). These elaborations feature a tricyclic fluorene moiety, a piperidine group, and a benzothiophene group (Figure 7). One molecule is an enantiomer of the other, suggesting that NCS-1 does not differentiate between them (Figure 7). While in Ric8inhib-3-125 the benzothiophene sulfur atom forms a H-bond with Y52, in Ric8inhib-3-21 the sulfur atom faces the opposite side of the cavity, establishing van der Waals with T92. Both elaborations are highly stabilized through π-π contacts with neighboring aromatic residues (Figure 7). An analysis of the interactions suggests that Ric8inhib-3-125 is a promising candidate for future development (Table 2).

Three structures derived from Ric8enhan-3 have been solved, all located in the middle region of the NCS-1 cavity (Figure 7). These elaborations share a common scaffold based on a sulfonamide, with a quinoline group on one side (Figure 7). The three structures solved show that the elaborations feature at the upper part of the cavity a benzene group with substituents at ortho-, meta-, and para-positions (Figure 7). Ric8enhan-3-666 is of particular interest, as it forms two H-bonds, with Y52 in the middle and with L89 at the top of the crevice. In addition, it has the largest contact area, establishing a higher number of interactions, which correlates with a better ΔG (Table 2). It is noteworthy that similar compounds containing an extra methyl group bound to the quinoline moiety were synthesized (Figure 5). None of these derivatives yielded structures, suggesting that the presence of the methyl group may destabilize the interaction with NCS-1.

**Table 2:**
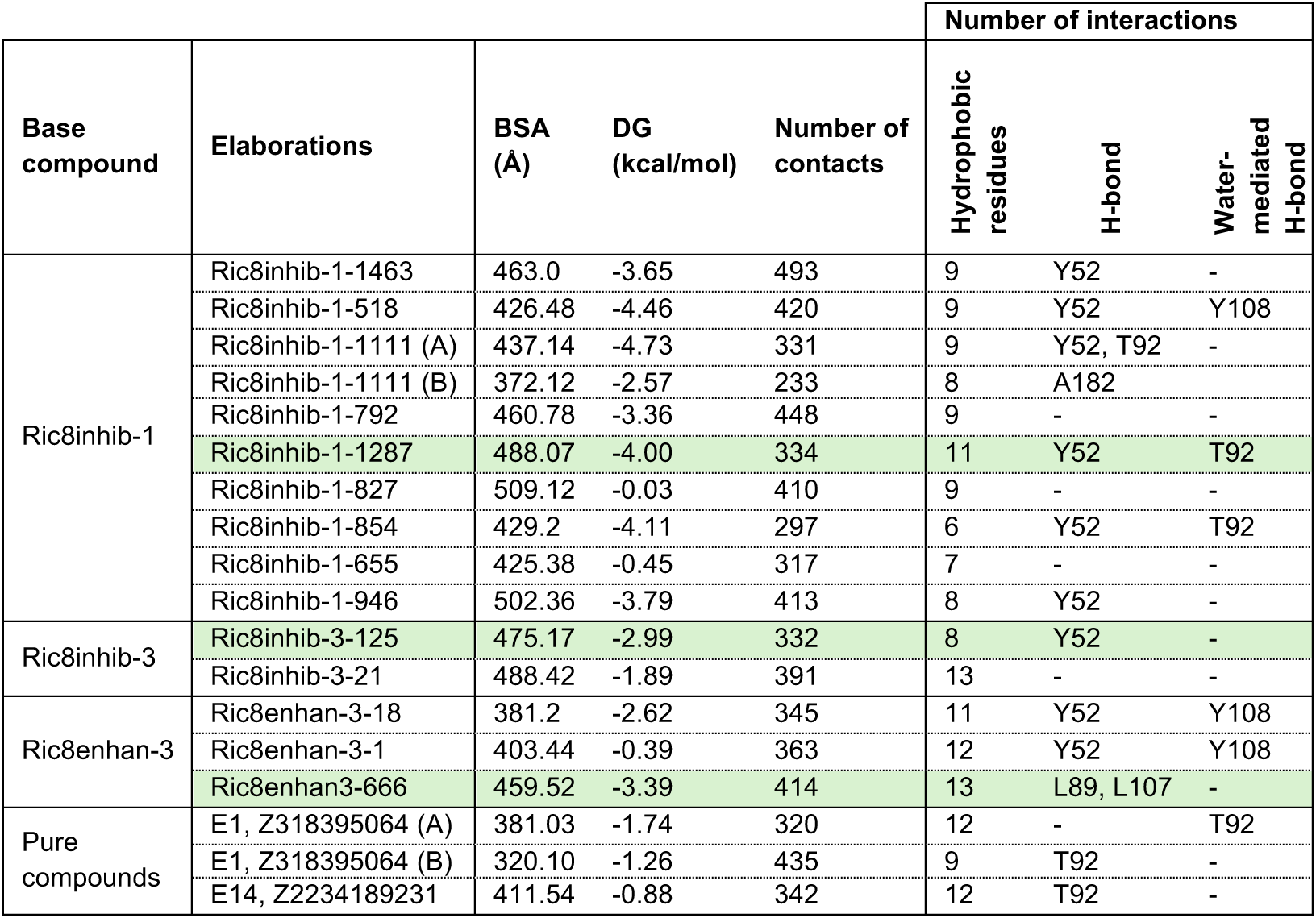
Analysis of the interactions found in the NCS-1 elaboration-bound structures. High-interest compounds are marked in green.

The structures of two commercially available pure compounds, E1 and E14, were solved with both bound at the upper half of the NCS-1 crevice, establishing H-bonds with residues in the middle part of the crevice and van der Waals contacts (Supplementary Figure 8 and Supplementary Table 2). In the case of E1, different binding modes were found when analyzing each independent NCS-1 molecule. Interestingly, E14 showed an affinity in the range of tens-to-hundred micromolar (Figure 6a and Supplementary Table 2), no affinity was detected for E1, which may be explained by the variable poses identified for this compound. Lastly, it is important to note that, although only two structures of the pure compounds were solved, these results come from a single crystal soaking experiment.

## DISCUSSION

### Implementation of the Fast Forward Fragment pipeline developed at XChem

To fully leverage structural information and efficiently explore the chemical space of biological targets, it is essential to develop workflows that enable rapid and cost-effective drug discovery. In this work, a fragment-based compound design approach has been used, applying an algorithmically guided fragment evolution and automated synthesis methodology to rapidly complete a DMTA cycle (Figure 2). The NCS-1 project represents the first project carried out at the XChem facility (DLS, Oxford, UK) where an iteration of the cycle has been completed, leading to the identification of compounds with micromolar affinities, which have the potential to regulate protein-protein interactions.

The crystallographic fragment screening has identified compounds with a wide chemical diversity and various interaction patterns with residues located in the NCS-1 crevice that are involved in target recognition (Figure 1 and 3)^7,11^,. The proximity of the fragments in the hydrophobic crevice of NCS-1 allowed fragment merging and linking (Supplementary Figure 6 and Supplementary Figure 7). As a result, six base compounds were chosen to potentially regulate the interaction of NCS-1 with Ric-8A, dopamine D_2_ and cannabinoid CB_1_ receptors, which served as inspiration to the rapid and cost-effective synthesis of 400 elaborations that broadly explored the PPI chemical space (Table 1, Figure 4 and 5). Furthermore, two semi-automated orthogonal methods were implemented to confirm the synthesis of the products. First, MSCheck has allowed the detection of synthetic products via LC-MS. Second, waveRAPID technology from the Creoptix system, has enabled the finding of NCS-1 binders, estimating affinities and kinetic constants of the interactions (Figure 6)^27^.

**Figure 8:**
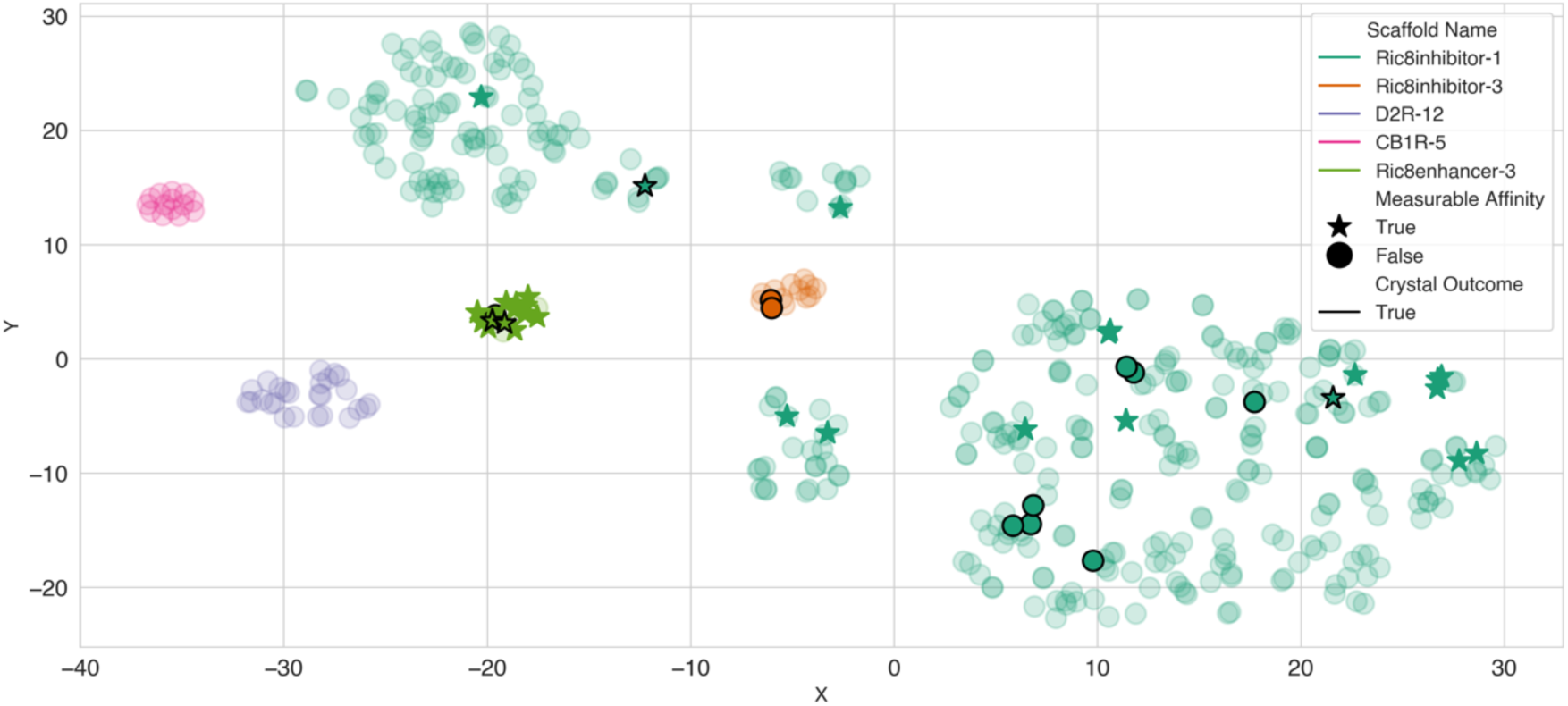
Exploration of the NCS-1 chemical space through synthetic elaborations. Chemical space of the 454 elaborations synthesized by CAR where the Morgan fingerprints (2048 bit length, radius=2) were reduced to two dimensions using the t-SNE algorithm. The color code is assigned to the different populations or clusters based on the common scaffold (base compound) they share: Ric8inhib-1 (teal), Ric8inhib-3 (orange), Ric8enhan-3 (light green), D2R-12 (purple), and CB1R-5 (pink). Compounds with at least one crystallographic structure are represented as dark circles with black outlines, compounds with measurable kinetics are represented as filled in stars, finally compounds with both crystallographic structures and measurable kinetics are stars with black outlines. Compounds with neither data are transparent circles.

While this experimental approach has yielded valuable structural insights, certain limitations should be acknowledged. First, although PanDDA has proven effective in many fragment screening campaigns, it is not universally applicable, particularly in systems with significant conformational flexibility like NCS-1. It’s also worth noting that newer versions, such as PanDDA2, may address some of these challenges. Second, structural divergence between larger compounds and their parent fragments has been observed, though this likely reflects the inherent complexity of the NCS-1 binding crevice rather than a limitation of the experimental method itself (Supplementary Figure 11, Supplementary Figure 12 and Supplementary Figure 13). Third, while many of the synthesized compounds are amide-based, a common motif in approved drugs, future optimization may still benefit from exploring more metabolically stable alternatives, depending on the intended pharmacokinetic profile^28–30^.

Finally, the completion of a full DMTA cycle has enabled the resolution of crystallographic structures of compounds that contribute to the exploration of the chemical space within the hydrophobic crevice of NCS-1 (Figures 7 and 8). The position of these compounds, along with the generated chemical diversity and interaction profiles (Figure 8), suggests that they may be selective against NCS-1 target proteins or display synergistic activities to different ones. It is worth highlighting that this strategy allowed us to obtain various scaffolds that differ structurally and chemically from previously obtained molecules such as FD-44, IGS-1.76, and 3b (Figure 1), which have not passed from preclinical stages in previous drug discovery programs^2,6,9^. Given the sequence homology and structural conservation of the NCS family of proteins, the fragments and larger compounds discovered might be a resource to generate new modulators for other pharmacologically relevant proteins of the family^5,7^.

In conclusion, this study has not only yielded promising compounds for further development as modulators of NCS-1 but also established a proof of concept for the novel methodologies developed and implemented at XChem. The results presented here will inform the refinement of underlying algorithms, contributing to the development of a robust and scalable workflow for the discovery of compounds with biomedical and biotechnological potential.

### Generation of lead compounds targeting NCS-1 PPI with therapeutic potential

NCS-1 is a pharmacological target of the central nervous system, which implies that modulators must cross the blood-brain barrier (BBB) and, therefore, should have a molecular mass below 400-500 Da^31^. Some fragments already exhibit molecular mass and interactions that suggest the feasibility of exploring vectorial growth strategies from a single fragment (Supplementary Figure 1b and Supplementary Figure 2b). Fragments located at the upper part of the NCS-1 crevice are particularly promising for modulating the interaction between NCS-1 and D_2_R in a specific manner, as the structural analysis of the protein-protein complex suggests. This is the case of fragment Z86416893 (Supplementary Figure 1b). On the other hand, fragment Z55290386, located in the central region of the NCS-1 crevice, establishes three hydrogen bonds with residues Y52 and T92, which are two hotspots of the NCS-1/Ric-8A interface (Supplementary Figure 2b). The positioning of this fragment suggests its potential for growth through vectors using two strategies: (i) growth toward the upper part of the cavity by inserting hydrophobic groups and (ii) growth toward the lower part of the cavity by inserting polar or hydrophobic groups, in order for example inhibit or stabilize the NCS-1/Ric-8A interaction^2,9^. In addition, fragment Z55290386 complies with Lipinski’s rule, indicating its compatibility as a potential oral drug. Moreover, according to prediction algorithms, this compound is predicted to cross the blood-brain barrier.

Three lead-like compounds are particularly interesting for the interaction patterns observed: Ric8inhib-1-1287, Ric8inhib-3-125, and Ric8enhan-3-666 (Table 2). These compounds could serve as starting points for the next round of optimization and further development. The predicted pharmacological properties of the 16 crystallized lead-like compounds suggest that none exhibit characteristics associated with problematic compounds that could form substructures flagged by PAINS alerts. Furthermore, all these compounds comply with Lipinski’s rule, indicating their potential as drug candidates. However, some molecules present Brenk alerts, which are associated with potential toxicity or chemical reactivity issues. These molecules are: Ric8inhib-1-1287, Ric8inhib-1-854, Ric8inhib-1111, Ric8enhan-3-1, and Ric8inhib-3-125. The BBB penetration prediction reveals that only Ric8inhib-3-21 is a candidate to cross this barrier, making it a potential therapeutic hit.

## METHODS

### Crystallization of NCS-1

Crystals of the full-length wild-type protein were obtained as previously reported^9^. The purified protein^9,32^ was dialyzed into a solution containing 20 mM sodium acetate pH 5.5, 0.5 mM CaCl_2_, and 0.5 mM DTT (2 changes, 4 and 16 h), and concentrated to 10 mg/ml (Vivaspin). Crystals grew in a crystallization condition containing 0.1 M sodium cacodylate trihydrate pH 6.5, 0.2 M sodium acetate trihydrate, 30% (v/v) polyethylene glycol 8000 (solution 28, Crystal Screen I, Hampton Research) at 4°C^9^. Crystallization was optimized using an Oryx robot (Douglas Instrument), refining the protein:precipitant ratio, to obtain higher diffraction quality and reproducibility. A final ratio of 200:300 nl produced larger and thicker plates in two days in 90% of the experiments. For fragment screening, crystallization experiments were conducted at the DCyBE (IQF-CSIC) while for lead-like compound screening, crystals were produced at XChem.

### High-throughput X-ray fragment screening

Four fragment libraries were selected: DSi-Poised (rapid progression and saturated heterocycles, 876 compounds)^33^, York 3D (substituted aliphatic heterocycles, 106 compounds)^34^, FragLites (halogenated, 31 compounds)^35^, and MiniFrags (Astex pharmacophores, 80 molecules)^36^, comprising a total of 1,093 different fragments. All of these were prepared in 100% DMSO at a concentration of 100 mM, 500 mM or 1 M. Crystallization plates were imaged using a Rock Imager (Formulatrix) at 4°C. The sites for dispensing the fragment-containing solution were selected with TexRank^37^, and site coordinates were recorded in SoakDB. Before screening, a DMSO tolerance test was conducted to assess crystal survival and diffraction quality. Incubation times ranged from 1-3 hours with solvent concentrations between 5-30%. After incubation, diffraction data were acquired. The optimal DMSO concentration and soaking time selected were 10% (v/v) and 2 hours at 4°C. Fragments were dispensed directly to the crystallization drops using the ECHO Liquid Handler (Labcyte), adjusted to a final DMSO concentration of 10% (v/v). Crystals were incubated for two hours at 4°C, then mounted on nylon loops (MiTeGen) using the Crystal Shifter (Oxford Lab Technologies), and flash-frozen in liquid nitrogen.

### Data collection and processing, structural resolution, refinement, and model validation

Diffraction data were collected at 100 K and 0.91507 Å wavelength at the beamline I04-1 (Diamond Light Source, UK)^12,38^. Large-scale analysis of protein-fragment complexes was performed with XChemExplorer (XCE)^39^. Diffraction data were automatically processed using AutoPROC^40^ and XIA2dials^41,42^. Initial map calculations were performed with Dimple ^43^. As search model, the previously solved NCS-1 structure from the same crystal form was used (PDB: 6QI4;^9^) with some modifications. Concretely, the dynamic N- and C-terminal ends were removed, and the search model only contained the region starting at EF-1 and ending at EF-4. A resolution limit of 3.5 Å was set to discard poorly diffracting datasets. Ligand restraint dictionaries were generated using aceDRG^44^. The ground-state model, representing an average model of the protein structure without a ligand, was built^45^ and used as a reference. An initial hit identification was performed with PanDDA^45^. Datasets with maps that showed additional density in the NCS-1 hydrophobic crevice or changes in the dynamic C-terminal helix H10 were selected for manual building to improve electron density maps and allow hit identification. This included remodeling of the N- and C-terminal ends of the protein. Ligands were modelled and bubble refinement focused on the fragment’s environment (10 Å distance) were carried out in Coot^46,47^. Structures were validated using MolProbity^48^ and Mogul^49^. The coordinates and structure factors were deposited in Fragalysis (fragalysis.diamond.ac.uk/) under the project code “NCS1”. All interactions were studied using LigPlot+^50^ and the PISA server^51^. The coordinates and structure factors have been deposited in the Protein Data Bank (PDB) under the group deposition ID G_1002352.

Data collection of fragment-based molecules was performed as described above. The structure of hNCS-1 (PDB: 6QI4) was used as search model and two refined structures, presenting different arrangements of the C-terminal helix H10 were used to solve the rest of the structures through Fourier differences calculations. Model building and refinement of all structures was performed with Coot and Phenix^46,52^. Supplementary File 1 and 2 show the statistics related to data processing and refinement.

### Progression of fragments: Linking and merging of fragments

Fragmenstein (https://github.com/oxpig/Fragmenstein)^20,21^ was used to elaborate the fragment-hits in a three-step pipeline: merging/linking, analogue-by-catalogue search, and placement. This algorithm is centered around strict obedience to the 3D position of the parent compounds, based on the premise that analogues with similar structures have a conserved binding mode, making ideal for crystallography-driven fragment-based drug discovery campaign. The workflow consisted of: (i) stitching together (link and merge) adjacent fragments or sub-fragments and energy minimizing in place with Fragmenstein via PyRosetta; (ii) searching for through catalogues using NextMove Software SmallWorld hosted at https://sw.docking.org^53^ to find commercially available analogs from Enamine, MCule and other suppliers within a graph edit distance of 5 from the merged compound; (iii) placing the candidate compounds with Fragmenstein and analysing the number of interactions, which are internally determined using the protein−ligand interaction profiler (PLIP)^54^. The virtual compounds were subset based on desired and undesirable interactions with key residues, and the ranked compounds filtered by eye. Due to dynamic nature of NCS-1, three templates were used. Additionally, some hypothesis-driven expansions were also enumerated using Next Move Software Arthor, hosted at https://arthor/docking.org^53^ and placed with Fragmenstein.

### Data dissemination and compound selection using Fragalysis

Fragalysis is a web application developed at XChem to facilitate structural data dissemination, provide GUI-based tools for data curation and annotation, and enable quality assessment by visually comparing the conformational and chemical features of in-silico follow-up designs (displayed on the right-hand-side) with experimental structural data (displayed on the left-hand-side). It also serves as a launch pad for structural biologists, computational chemists and medicinal chemists to collaborate, via shareable web links, with the ultimate endpoint of deciding how project resources should be allocated to progress fragments. In this work, NCS-1 structural data were uploaded to Fragalysis, key fragments were identified and subsequently used as inputs, with the experimental data standardized in Fragalysis, for computational efforts and compound merging via Fragmentstein. Critically, Fragalysis was used by the project team to review and select designs for further elaboration via the Syndirella and HIPPO workflow, for final building block purchase and CAR synthesis.

### Automated design and synthesis of new compounds using CAR

#### Compound design and selection for synthesis via CAR

Base compounds for further elaboration through digitized chemical synthesis were selected using Syndirella (https://github.com/kate-fie/syndirella). This algorithmic tool produces elaborations from the base compounds that are restricted to specific synthetic routes. First, a retrosynthesis analysis was performed using the tool Manifold (PostEra Inc., https://app.postera.ai/manifold/). Then a catalogue search for superstructures (molecules which contain the query molecule as a maximum common substructure) of each reactant was performed using Manifold. The catalogues selected were Enamine BB, Enamine REAL, Enamine MADE^55^, MCule and MCule Ultimate. Elaborated products were produced by performing a cartesian combination of reactants and reacted together *in-silico* with SMARTS patterns. Finally, the elaborations were filtered using Fragmenstein, to discard elaborations that would result in steric clashes with the protein after minimization to the inspiration fragments. Only elaborations using single step synthesis routes were selected. Syndirella elaborations were verified to have <0 kcal/mol ΔΔG_bind_ values and <1 Å RMSDs to the inspiration fragments, showing the selected mergers share a significant three-dimensional space with their inspiration fragments (Table 1). However, the Tanimoto similarity coefficient was relatively low, indicating the introduction of atoms or chemical groups different from those in the inspiration fragments. Despite the low coefficient, the mergers were expected to recapitulate the interactions of the inspiration fragments (Table 1, Supplementary Figure 6 and Supplementary Figure 7).

#### Classification of compounds for potential PPI modulation of NCS-1 with target proteins

Key amino acids involved in the NCS-1 PPI with D_2_R, Ric-8A and CB_1_R were identified and analyzed. For potential NCS-1/ D_2_R modulators, hydrophobic contacts with residues W30, L89 and I51 were considered, as well as D37, Y52 and Q181 or their ability to establish H-bonds^11^. For the NCS-1/Ric-8A interaction, classification criteria included hydrophobic contacts with W30 and hydrogen bonds with Y52 and T92 in the middle part of the crevice^7^. Additional interactions with the C-terminal helix H10 were also evaluated to identify potential Ric-8A inhibitors as previously reported^2,6^. For finding Ric-8A stabilizers, polar groups introduced in the middle of the crevice were considered essential for enhancing the PPI ^9^. Furthermore, an AlphaFold Multimer model of the NCS-1/CB_1_R complex, based on experimental results, indicated that residues Y52, Y108, R148, and R151 may be involved in CB_1_R recognition. These residues were therefore identified as critical for modulating the NCS-1/ CB_1_R interaction.

#### Starting material collections

HIPPO algorithm (Hit Interaction Profiling for Procurement Optimisation) (https://github.com/mwinokan/HIPPO) was used to select the best collection of elaborated products to maximize the use of reagents, synthesizing the highest number of compounds with the minimum number of reagents. Starting material combinations were also considered to increase chemical diversity and explore new interactions. All starting materials were sourced through MCule.

#### Automated Robotic Synthesis

Automated synthesis of the compounds was performed using CAR (Chemist Assisted Robotics), a digitized and database driven chemistry platform that facilitates multi-step high-throughput synthesis. The process involved: (i) Reviewing the synthetic routes proposed by Syndirella; (ii) automated generation of Python scripts and starting materials plate preparation for synthesis using an OpenTrons II liquid-handling robot; (iii) the execution of the chemical synthesis using an OpenTrons OT-2 robot. Control reactions validated the generated scripts. Starting materials (building blocks acquired from MCule) were dissolved in 100% DMF at a final concentration of 0.5 M and reactions were designed to yield the final product at a 100 mM concentration. Reactions were carried out in a 384-ECHO-LDV plate at a final volume of 10 µl, within a fume hood at 23°C. For amidation reactions, HATU was used as an activator and DIPEA as the base. Amidation reactions were performed in three different batches. Sulfonamidation reactions (Schotten-Baumann reactions with amines) used DIPEA as the base and were performed in a single batch. The reaction mixtures were sonicated for 12 hours at room temperature with agitation. Post-reaction, the mixtures were concentrated using SPE-Dry and solvent exchanged from DMF to DMSO by drying the ECHO plate for 30 minutes, then adding 10 µl of DMSO to reach a final concentration of 100 mM. 454 reactions were completed over four days.

#### Product analysis via LC-MS

0.5 µl of each reaction mixture was dissolved in 100 µl of acetonitrile. LC-MS data were collected using an Agilent 1260 HPLC system coupled with an electrospray ionization mass spectrometer G6120 LC-MS Single Quad and a Supelco Ascentis C18 column. The mobile phase consisted of 0.1% formic acid in water (Phase A) and acetonitrile (Phase B). Synthesized products were identified using MSCheck^24^, a semi-automated LC-MS data analysis tool that detects M+H^+^ and M+Na^+^ ion pairs from chromatogram peaks, where M is the molecular mass of the synthesis product. A manual identification of smaller synthesis peaks was also conducted to detect lower conversion rates. The final products were not purified.

### SAR by catalog

In addition to CAR synthesis, compounds from commercial catalogs were selected to expand the chemical diversity of mergers obtained through Fragmenstein. These compounds were purchased from Enamine (Supplementary Table 1).

### High-throughput screening of crude reaction mixtures and pure compounds via X-ray crystallography

As with the fragments, incubations and diffraction data collection were performed on synthesized compounds using crude reaction mixtures from CAR and pure available compounds. An initial round of screening involved all synthesized compounds with 2-hour incubations. A second round focused only on synthesis products detected via LC-MS, using incubation times of 8 and 18 hours per reaction mixture.

### Screening of compounds in solution via grating-coupled interferometry (GCI)

The Creoptix WAVEdelta system (Creoptix AG) equipment was used to measure binding affinities. Data analysis was carried out using software integrated with the equipment, which performed corrections based on DMSO calibration curves, using blank injections (FC1) as a reference. For pure compounds, kinetic parameters R_max_, k_on_, k_off_, and K_d_ were automatically calculated by the software through fitting to a 1:1 equilibrium model, using the RAPID assay, which depends on the ligand concentration. For reaction mixtures, an adjustment was made based on the dissociation rate constant (k_off_), as the exact concentration of products in the reaction mixtures is unknown^16^. Evaluation criteria include a R_max_ limit (15 pg/mm^2^) based on the control E17 compound, the binding profile of samples on both active and blank channels and checking the non-specific binding to the blank channel. After the fitting, the dissociation curves and the k_d_ values were reviewed to exclude the false positive hits. The off-rate screening assay was performed in duplicate, and the binders were chosen based on the binding response and k_d_ values in both runs. The control E17 sample was repeatedly included in the assay to monitor the NCS-1 activity and binding response over the run.

### Tryptophan emission fluorescence

NCS-1 emits intrinsic fluorescence when excited at 295 nm, due to the presence of Trp and Tyr residues located at the hydrophobic crevice. The binding of compounds to the crevice changes the 3D environment of these residues and allows the estimation of an apparent affinity constant^5^. The fluorescence intensity was measured at 330 nm in a nano-DSF apparatus (nanoTemper) at increasing concentrations of the pure compounds at 35°C. Protein concentration was set to 5 μM and ligands, from 25 μM to 250 μM in a final buffer containing 50 mM Tris pH 8.0, 125 mM NaCl, 5 μM CaCl_2_ and 5% DMSO. Three independent experiments were performed with each compound. The apparent dissociation constant, K_d_ was calculated using a least squares algorithm to fit the recorded data to the equation previously described^5^.

### Analysis of interactions between NCS-1 and fragment-based molecules

Interactions were analyzed using NCONT from CCP4 with a 4.2 Å interaction limit cut-off^56^, LigPlot+^50^ and the PISA server^51^. ADME Tox (Absorption, Distribution, Metabolism, Excretion, and Toxicity) properties were estimated using SwissADME^57^.

## ACKNOWLEDGMENTS

M.J.S.-B. would like to thank the following funding agencies: “Agencia Estatal de Investigación”-Spanish Ministry of Science, Innovation and Universities (AEU-MICIU grants PID2022-137331OB-C31/ AEI/10.13039/501100011033/ FEDER, UE) and the Spanish National Research Council (2023AEP005 and iMOVE-23223 to D.M.R.). Access to Diamond Light Source and XChem facility was granted by BAG Proposal No. Ib31306, Instruct-ERIC (2023 APPID 27355) and iNEXT-Discovery (PID24398). D.M.R. was funded with an Instruct-ERIC Internship (13th Instruct Call, APPID 3080) and C.M.R. with a CSIC JAE-Intro (JAEINT_23_00101). M.J.S.B. would like to thank Dr. Inmaculada Pérez Dorado, coordinator of the DLS-XChem BAG for her support. A.M. is funded by “Agencia Estatal de Investigación”-Spanish Ministry of Science (grant PID2022-137331OB-C32). K.F. was supported by funding from the Medical Sciences Doctoral Training Centre, University of Oxford and IBM Research.

## DATA AVAILABILITY

The atomic coordinates and structure factors of the NCS-1 in complex with the ligands have been deposited in the Protein Data Bank, https://www.pdb.org/. Structures containing fragment compounds are available under the group deposition ID G_1002352, with accession codes: 7IK9, 7IKA, 7IKB, 7IKC, 7IKD, 7IKE, 7IKF, 7IKG, 7IKH, 7IKI, 7IKJ, 7IKK, 7IKL, 7IKM, 7IKN, 7IKO, 7IKP, 7IKQ, 7IKR, 7IKS, 7IKT, 7IKU, 7IKV, 7IKW, 7IKX, 7IKY, 7IKZ, 7IK0, 7IK1, 7IK2, 7IK3, 7IK4, 7IK5, 7IK6, 7IK7, 7IK8, 7IK9, 7ILA, 7ILB, 7ILC, 7ILD, 7ILE, 7ILF, 7ILG, 7ILH, 7ILI, 7ILJ, 7ILK, 7ILL. Accession codes for structures containing fragment-based larger compounds are: 9QK7, 9QK8, 9QK9, 9QKA, 9QKB, 9QKC, 9QKD, 9QKE, 9QKF, 9QKG, 9QKH, 9QKI, 9QKJ, 9QKK, 9QKL and 9QKM. Authors will release the atomic coordinates upon article publication.

## AUTHOR INFORMATION

The authors declare no conflict of interest.

## Author contributions

D.M.R.: investigation, data analysis, writing original draft, reviewing and editing. E.C.: investigation, data analysis and writing. K.K.F., M.W. and M.F.: data analysis, writing. S.P.-S., C.M.R., A.M. and M.G.: investigation, data analysis. P.G.M. and C.W.E.T: data collection. W.T. and D.F.: conceptualization, investigation, data analysis, writing and reviewing. F.vD.: conceptualization, reviewing, resources, funding acquisition, validation, supervision, and project administration. M.J.S.B.: conceptualization, investigation, data analysis, writing original draft, reviewing and editing, resources, funding acquisition, validation, supervision, and project administration.

