## Supplementary Figures and Tables for "Combining crystallographic fragment screening, algorithmic merging and digitized chemical synthesis efficiently yields scaffolds that target diverse NCS-1 protein-protein interactions implicated in neurological disorders"

[1]Department of Crystallography and Structural Biology. Institute of Physical-Chemistry "Blas Cabrera", CSIC, Serrano 119, Madrid 28006, Spain.
[2]Department of Statistics, University of Oxford, Oxford OX1 3LB, United Kingdom.
[3]Research Complex at Harwell, Harwell Science and Innovation Campus, OX11 0FA, Didcot, United Kingdom.
[4]Centre for Medicines Discovery, University of Oxford, Oxford OX3 7DQ, United Kingdom.
[5]Diamond Light Source Ltd, Harwell Science and Innovation Campus, OX11 0QX, Didcot, United Kingdom.
[6]Department of Neurobiology, Instituto Ramón y Cajal de Investigación Sanitaria, Hospital Universitario Ramón y Cajal, Madrid (Spain).
[7]Department of Biochemistry, University of Johannesburg, Johannesburg 2006, South Africa.

### List of supplementary material

**Supplementary figures:**

**Figure S1:** Fragments binding to the upper region of the NCS-1 cavity (Sites 1 and 2).
**Figure S2:** Fragments binding to the middle region of the NCS-1 cavity.
**Figure S3:** Fragments binding to the lower region of the NCS-1 cavity (Sites 4-6).
**Figure S4:** Fragments binding to the lower region of the NCS-1 cavity (Sites 7-10).
**Figure S5:** Predicted interactions of pure compounds.
**Figure S6:** Base compounds selected as potential modulators of the interaction between NCS-1 and dopamine $D_2$ or $CB_1$ receptors for synthetic elaboration via CAR.
**Figure S7:** Base compounds selected as potential modulators of the interaction between NCS-1 and Ric-8A for synthetic elaboration via CAR.
**Figure S8:** Overview of the crystallographic structures of the bigger compounds.
**Figure S9:** Structures of elaborations derived from fragments.
**Figure S10:** The binding of larger, commercially available compounds to NCS-1 using tryptophan emission fluorescence.
**Figure S11:** Superimposition of crystallographic structures of Ric8inhib-1 elaborations with poses predicted by Fragmenstein.

**Figure S12:** Superimposition of crystallographic structures of various elaborations relative to placements predicted by Fragmenstein.

**Figure S13:** Superimposition of crystallographic structures of various elaborations relative to crystallographic structures of their inspiration fragments.

**Supplementary tables:**

**Table S1:** Selected pure catalog compounds.

**Table S2:** Comparison of affinity results obtained via GCI and Trp fluorescence emission techniques.

**Table S3:** Summary of the different steps during GCI for NCS-1 capture on the NTA surface.

### Supplementary figures

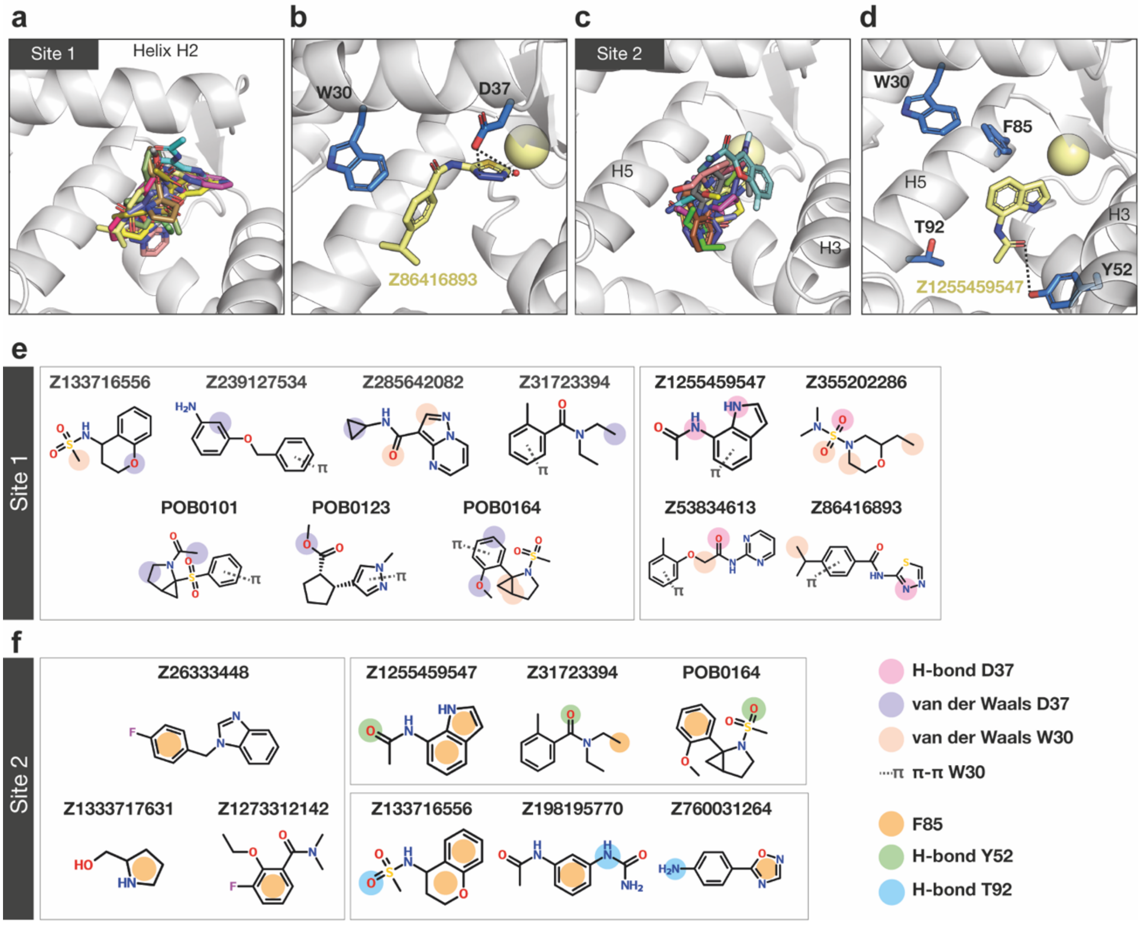


**Figure S1: Fragments binding to the upper region of the NCS-1 cavity (Sites 1 and 2).** (**a**) Superposition of fragments interacting with helix H2 (Site 1). NCS-1 is shown in ribbon form, and the fragments are displayed as sticks in different colors. (**b**) Representative figure of the interactions of the fragments at Site 1. Only key residues, W30 and D37, are shown as blue sticks, while the fragment Z86416893 is displayed as yellow sticks. (**c**) Superposition of fragments interacting with F85 and residues from helices H3 and H5 (Site 2). (**d**) Representative figure of the interactions of fragments binding at Site 2. Key interacting residues W30, Y52, F85, and T92 are displayed. (**e**) and (**f**) 2D structures of unique fragments and their interactions with NCS-1 at Site 1 (E) and Site 2 (F). Hydrogen bonds with D37 (pink), van der Waals interactions with D37 (light purple), van der Waals interactions with W30 (light orange), and π-π interactions with W30 are indicated. Additionally, hydrophobic and π-π interactions with F85 (orange), hydrogen bonds with Y52 (green), and hydrogen bonds with T92 (light blue) are represented.

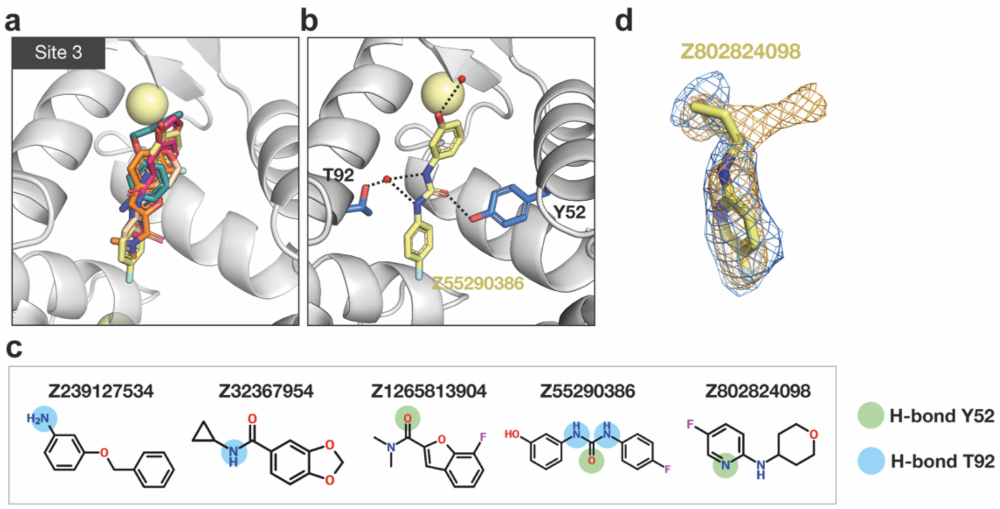


**Figure S2: Fragments binding to the middle region of the NCS-1 cavity.** (**a**) Superposition of fragments located at Site 3. (**b**) Representative figure of the interactions occurring at Site 3. The fragment Z55290386 forms hydrogen bonds with Y42 and T92. (**c**) 2D structures of unique fragments at Site 3. Hydrogen bonds with Y52 (green) and T92 (blue) are represented. (**d**) Despite some ambiguity in the electron density around the tetrahydropyran ring of fragment Z802824098, the binding pose of the pyridine ring provided valuable insight for follow-up design. The PanDDA event map (orange) suggested two possible conformations for the tetrahydropyran ring, both of which were initially modeled and refined. However, the 2Fo–Fc map (blue) indicated that one conformation was predominant, leading to the removal of the low-occupancy alternative.

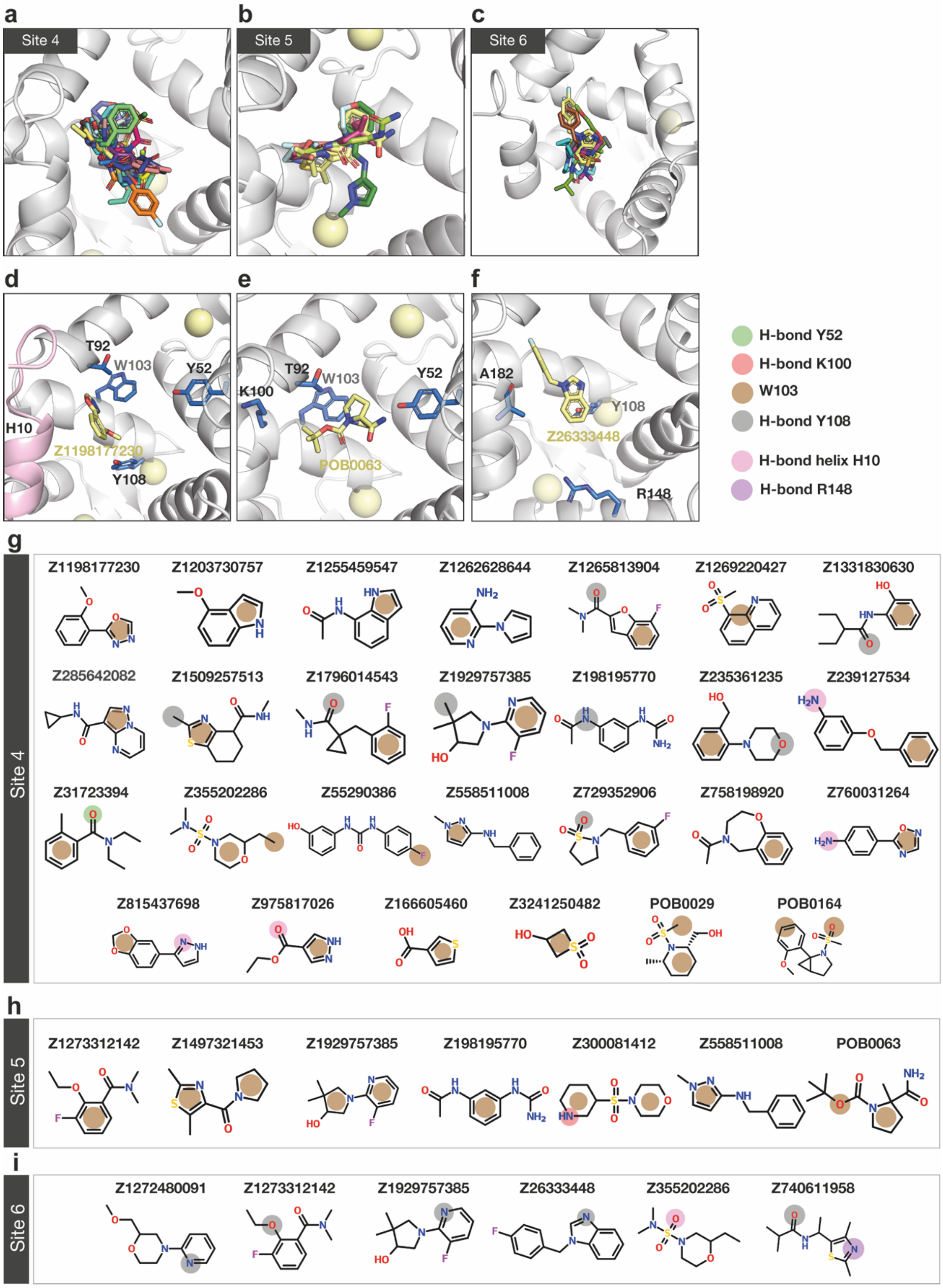


**Figure S3: Fragments binding to the lower region of the NCS-1 cavity (Sites 4-6).** (**a**), (**b**), (**c**) Superposition of fragments binding at Sites 4, 5, and 6, respectively. (**d**), (**e**), (**f**) Representative figures of the interactions occurring at Sites 4, 5, and 6, respectively. Fragments Z1198177230 (Site 4), POB0063 (Site 5), and Z26333448 (Site 6) are displayed as yellow sticks. The relevant NCS-1 side chains involved in recognition are shown as blue sticks. (**g**), (**h**), (**i**) 2D structures of the fragments and interactions with NCS-1. Hydrogen bonds with Y52 (green), K100 (red), Y108 (gray), residues from helix

H10 (pink), and R148 (purple) are represented. Additionally, van der Waals contacts with W103 (brown) are shown.

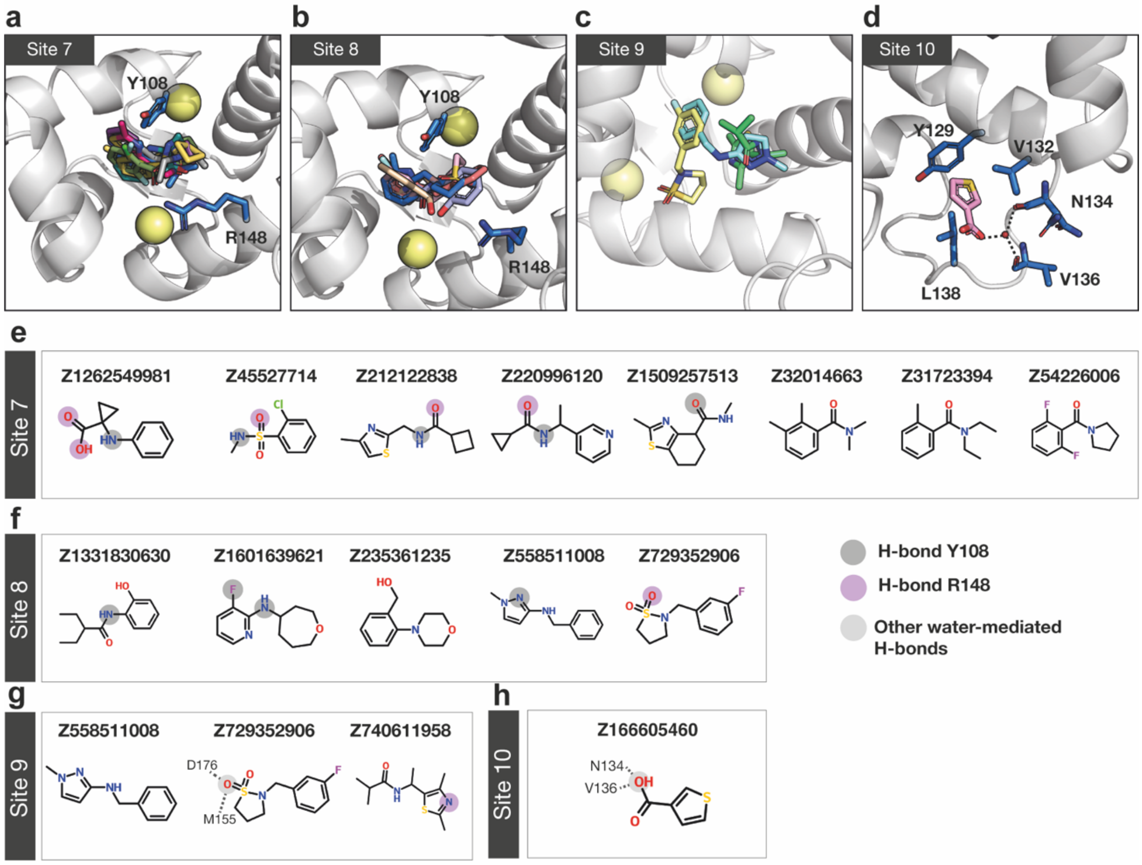


**Figure S4: Fragments binding to the lower region of the NCS-1 cavity (Sites 7-10).** (**a**), (**b**), (**c**), (**d**) Superposition of fragments binding at Sites 7, 8, 9, and 10, respectively. (**e**), (**f**), (**g**), (**h**) 2D structures of the unique fragments. Only hydrogen bonds with Y108 (gray), R148 (purple), and other bonds mediated by water molecules (light gray) are represented.

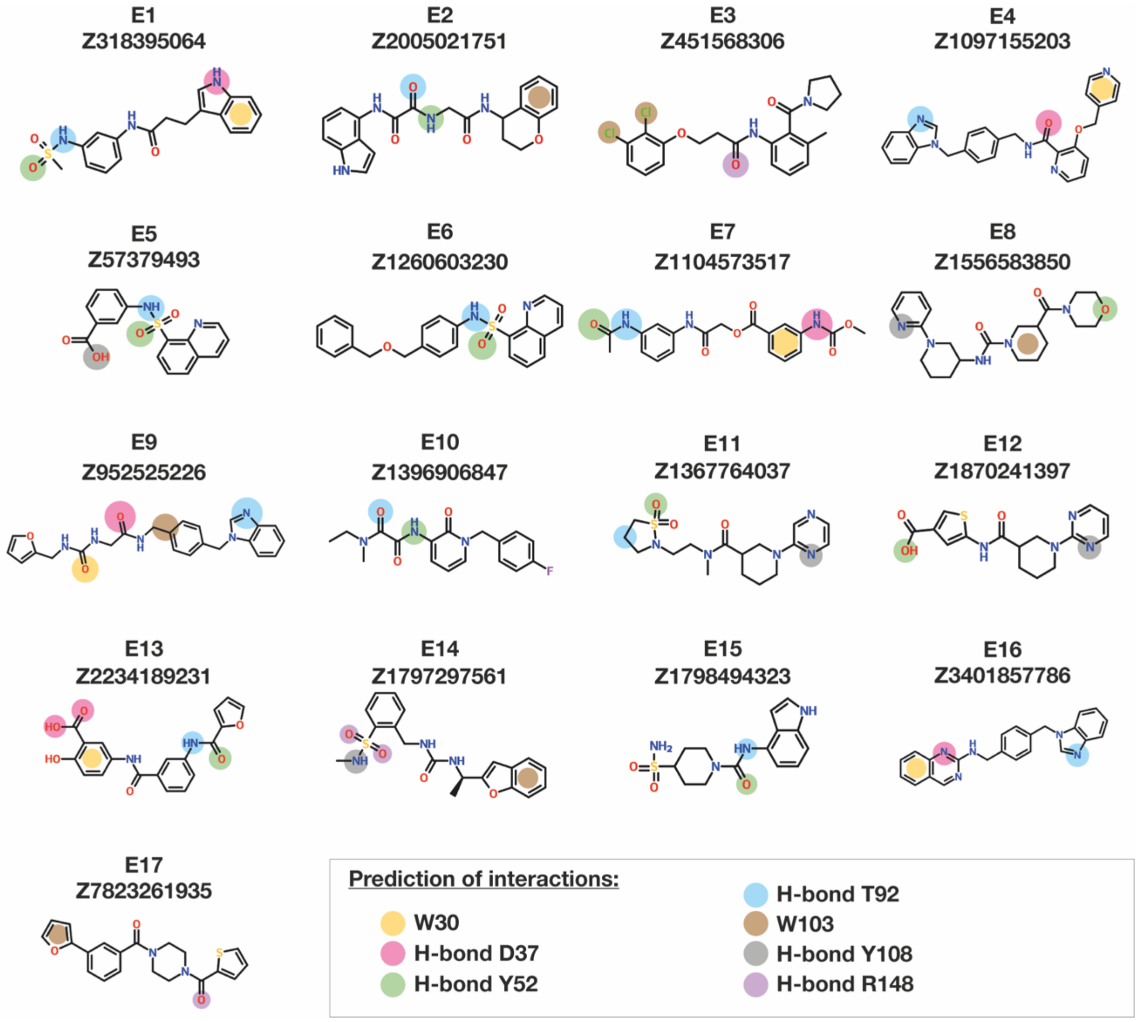


**Figure S5: Predicted interactions of pure compounds.** The 2D structure of each compound is shown, along with the assigned code (E1-E17) and the commercial code from Enamine. The predicted interactions are represented by circles of different colors.

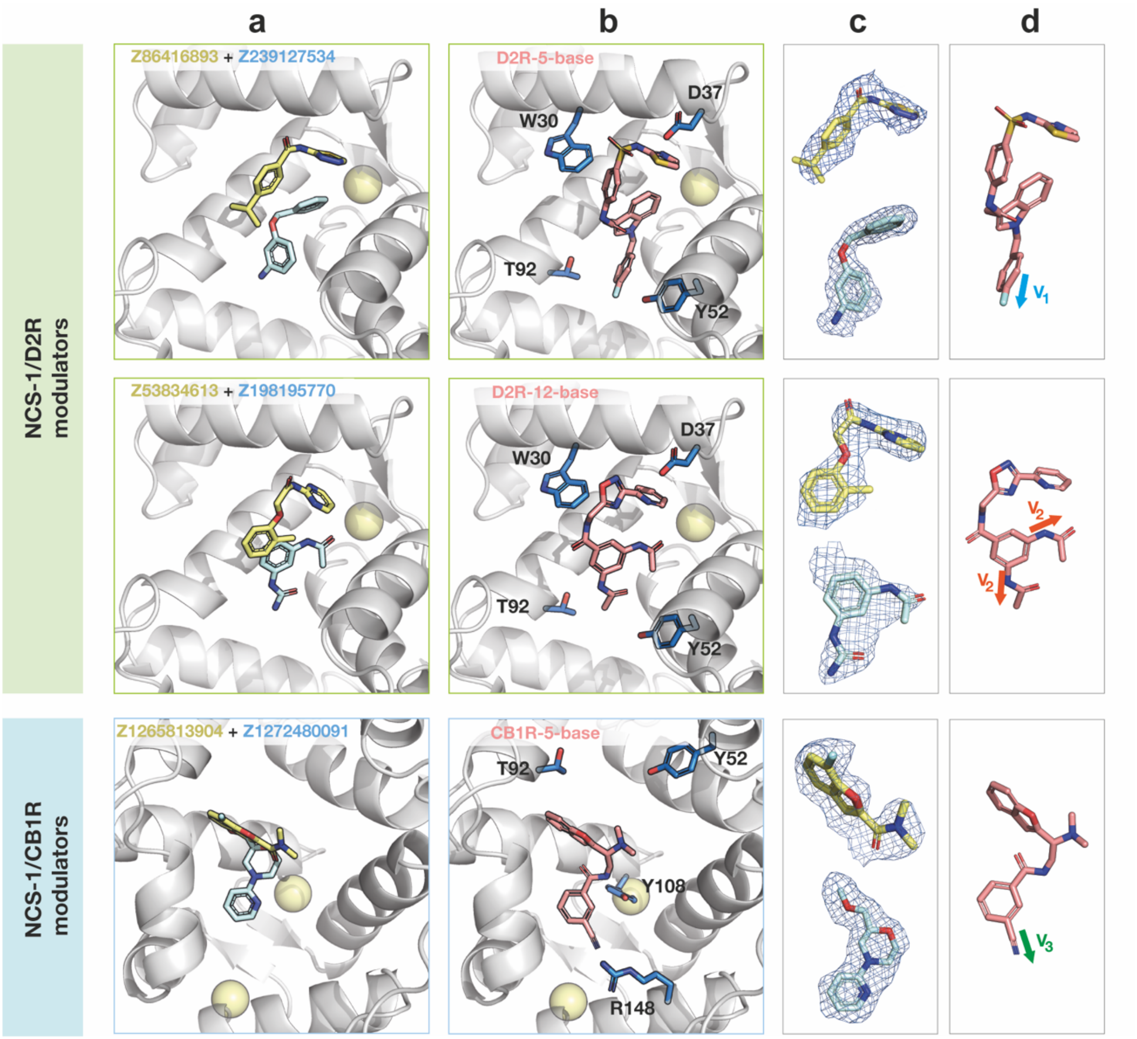


**Figure S6: Base compounds selected as potential modulators of the interaction between NCS-1 and dopamine $D_2$ or $CB_1$ receptors for synthetic elaboration via CAR.** (**a**) The crystallographic structure of the fragments (blue and yellow) that inspired the creation of the merged compounds is shown. (**b**) Representation of the placement using Fragmenstein. The key NCS-1 residues (in dark blue) and the theoretical three-dimensional structure of the mergers (in salmon color) are shown. (**c**) Electron density map 2Fo-Fc of the inspiration fragments (contour at 1.0 σ). (**d**) Vectors (Vx) detected in the theoretical structures of the base compounds for molecules elaborations.

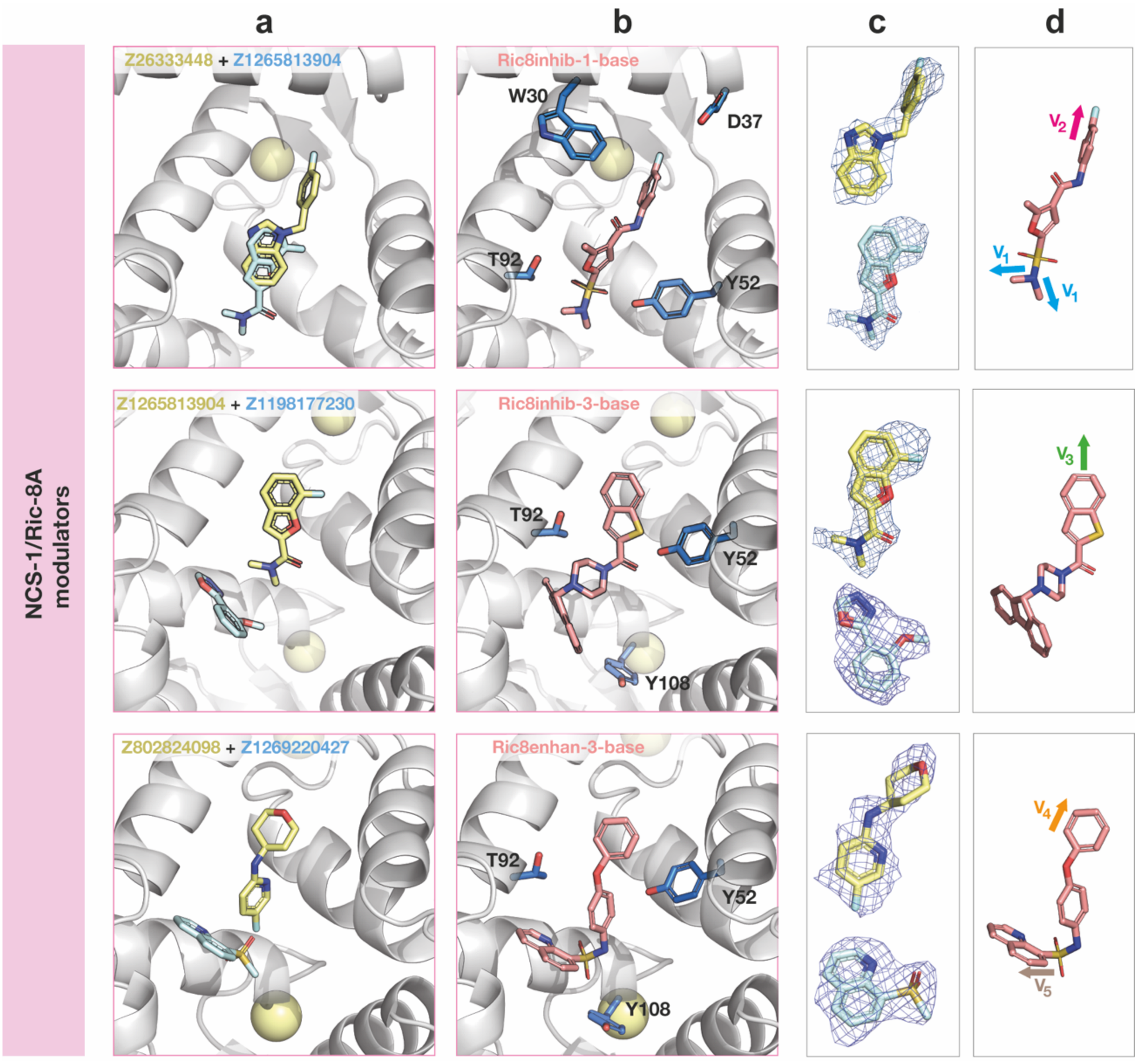


**Figure S7: Base compounds selected as potential modulators of the interaction between NCS-1 and Ric-8A for synthetic elaboration via CAR.** (**a**) Crystallographic structure of the inspiration fragments. Fragments are shown as yellow and blue sticks. (**b**) Placement representation using Fragmenstein in the NCS-1 crevice (theoretical structures). The key interaction residues and the theoretical three-dimensional structure of the merger (salmon) are shown. (**c**) Electron density map 2Fo-Fc of the inspiration fragments (contour at 1.0 σ) in dark blue. (**d**) Visual vectors detected in the theoretical structures of the base compounds for progression via elaboration.

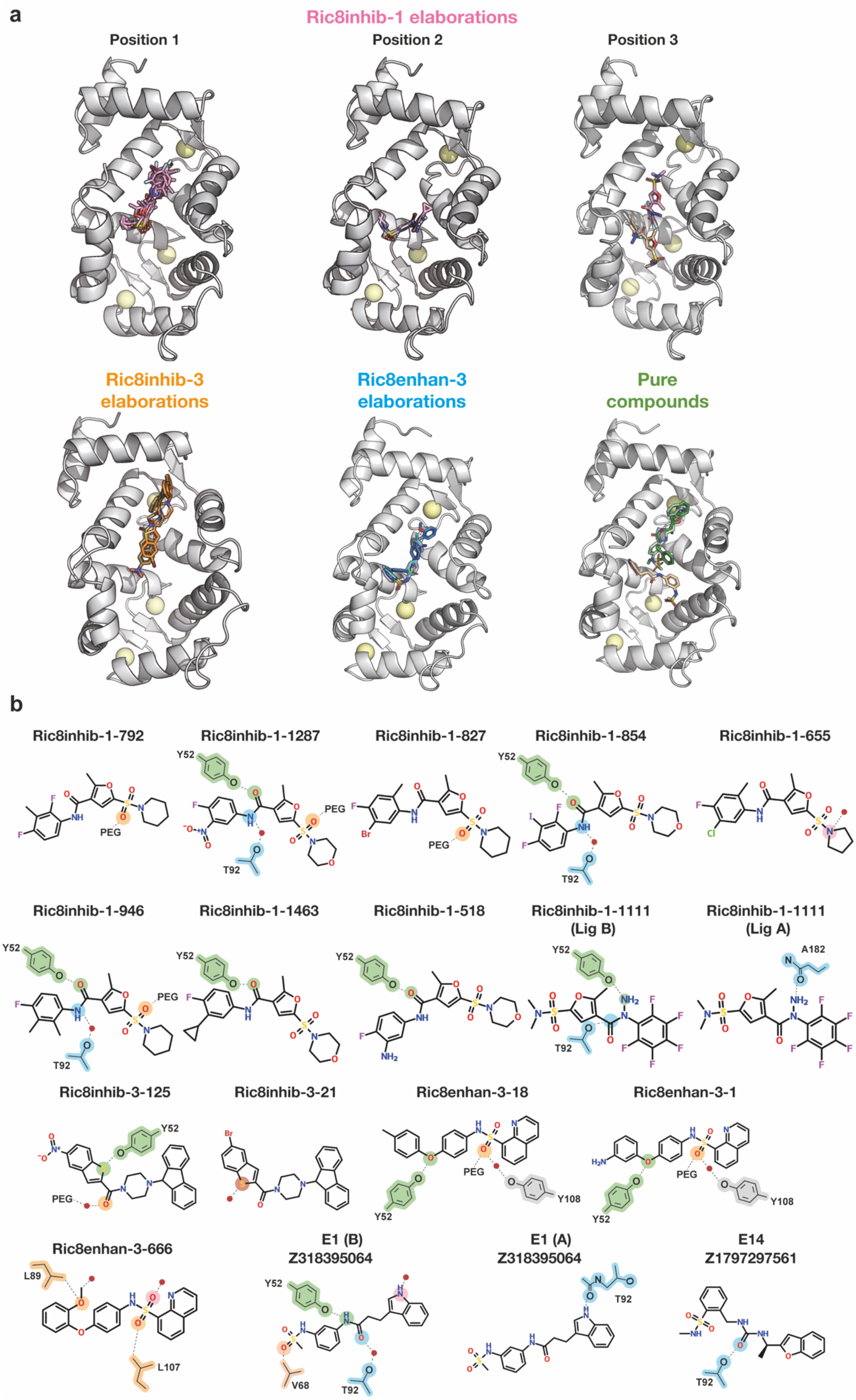
a
Ric8inhib-1 elaborations
Position 1
Position 2
Position 3
Ric8inhib-3 elaborations
Ric8enhan-3 elaborations
Pure compounds
b
Ric8inhib-1-792
Ric8inhib-1-1287
Ric8inhib-1-827
Ric8inhib-1-854
Ric8inhib-1-655
Ric8inhib-1-946
Ric8inhib-1-1463
Ric8inhib-1-518
Ric8inhib-1-1111 (Lig B)
Ric8inhib-1-1111 (Lig A)
Ric8inhib-3-125
Ric8inhib-3-21
Ric8enhan-3-18
Ric8enhan-3-1
Ric8enhan-3-666
E1 (B) Z318395064
E1 (A) Z318395064
E14 Z1797297561
Y52
T92
PEG
A182
Y108
L89
L107
V68

**Figure S8: Overview of the crystallographic structures of the bigger compounds.** (**a**) Superposition of the different crystallized elaborations related to base compounds Ric8inhib-1 (pink), which have been classified in three different positions (Position 1, 2, and 3), Ric8inhib-3 (orange), Ric8enhan-3 (blue) and pure Enamine compounds (green). $Ca^{2+}$ ions are depicted as yellow spheres. (**b**) 2D structure of the crystallized elaborations showing H-bond interactions (dashed line) with NCS-1.

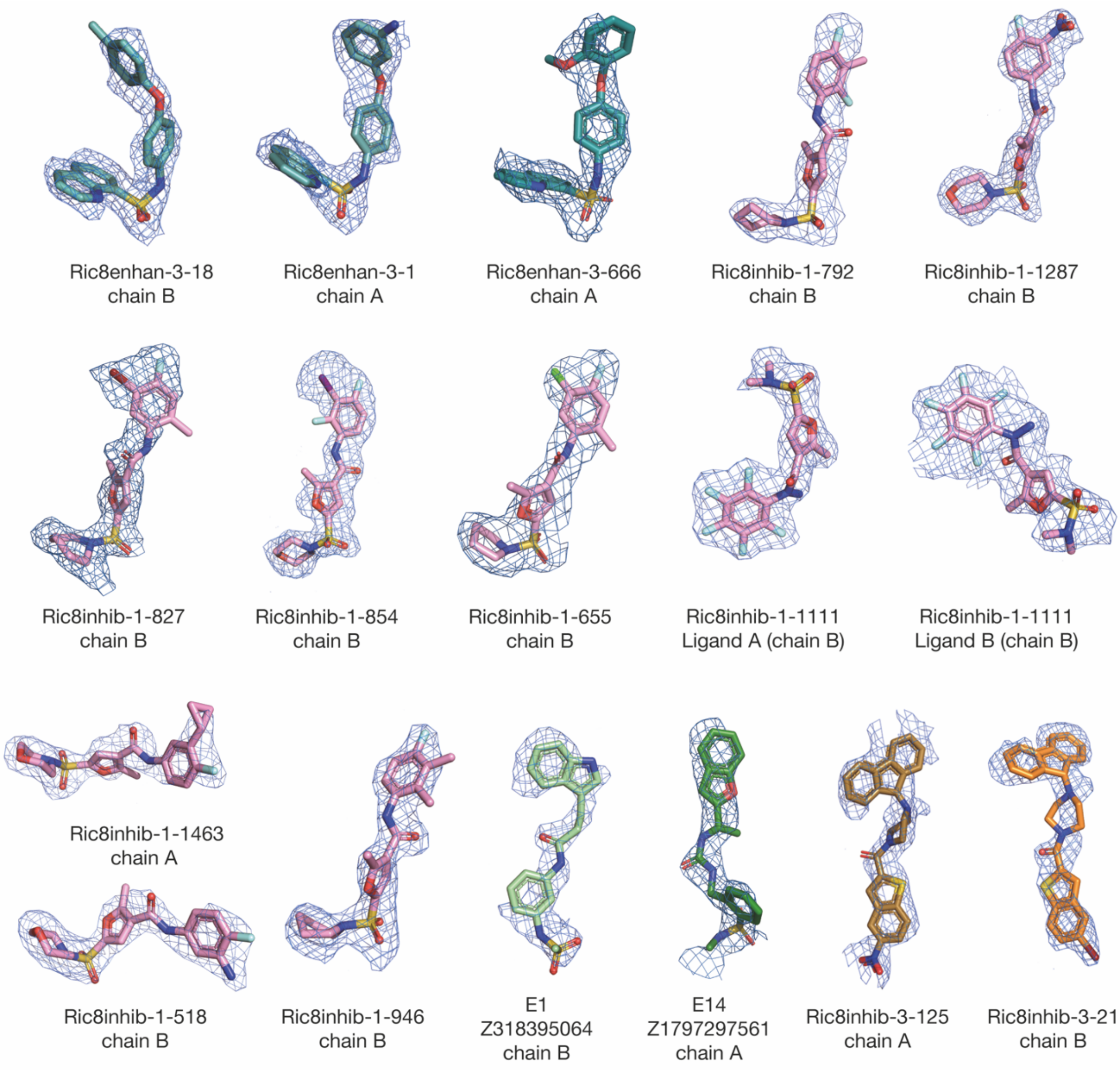


**Figure S9: Structures of elaborations derived from fragments.** The 2Fo-Fc electron density map is shown in dark blue, contoured at 1σ. The Ric8inhib-3-125 map is contoured at 0.8σ. The protein chain where the ligand is located is indicated (chain A or B). 2 molecules of Ric8inhib-1-1111 bind to the NCS-1 crevice, as specified.

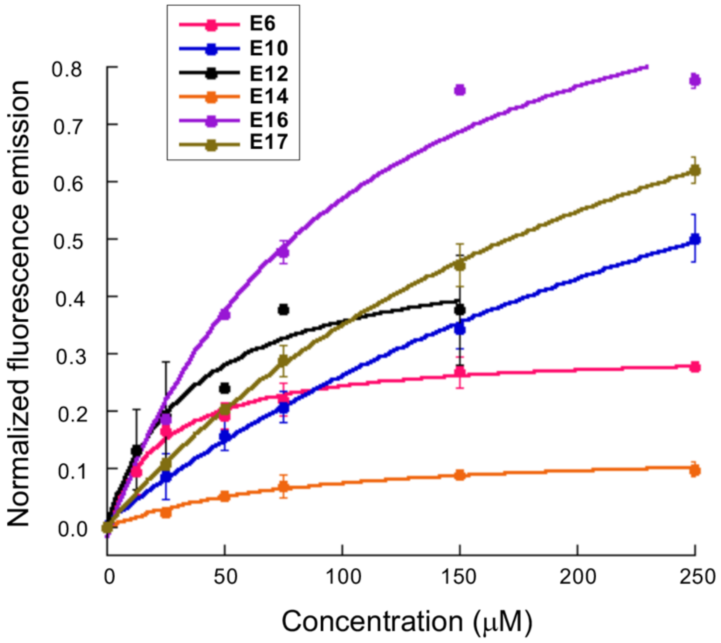


**Figure S10: The binding of larger, commercially available compounds to NCS-1 using tryptophan emission fluorescence.** Normalized intrinsic fluorescence emission of NCS-1 (mean ± SEM, n=3) at increasing concentrations of the pure compounds. Curves correspond to the least squares fitting of the recorded data to a 1:1 stoichiometry equilibrium.

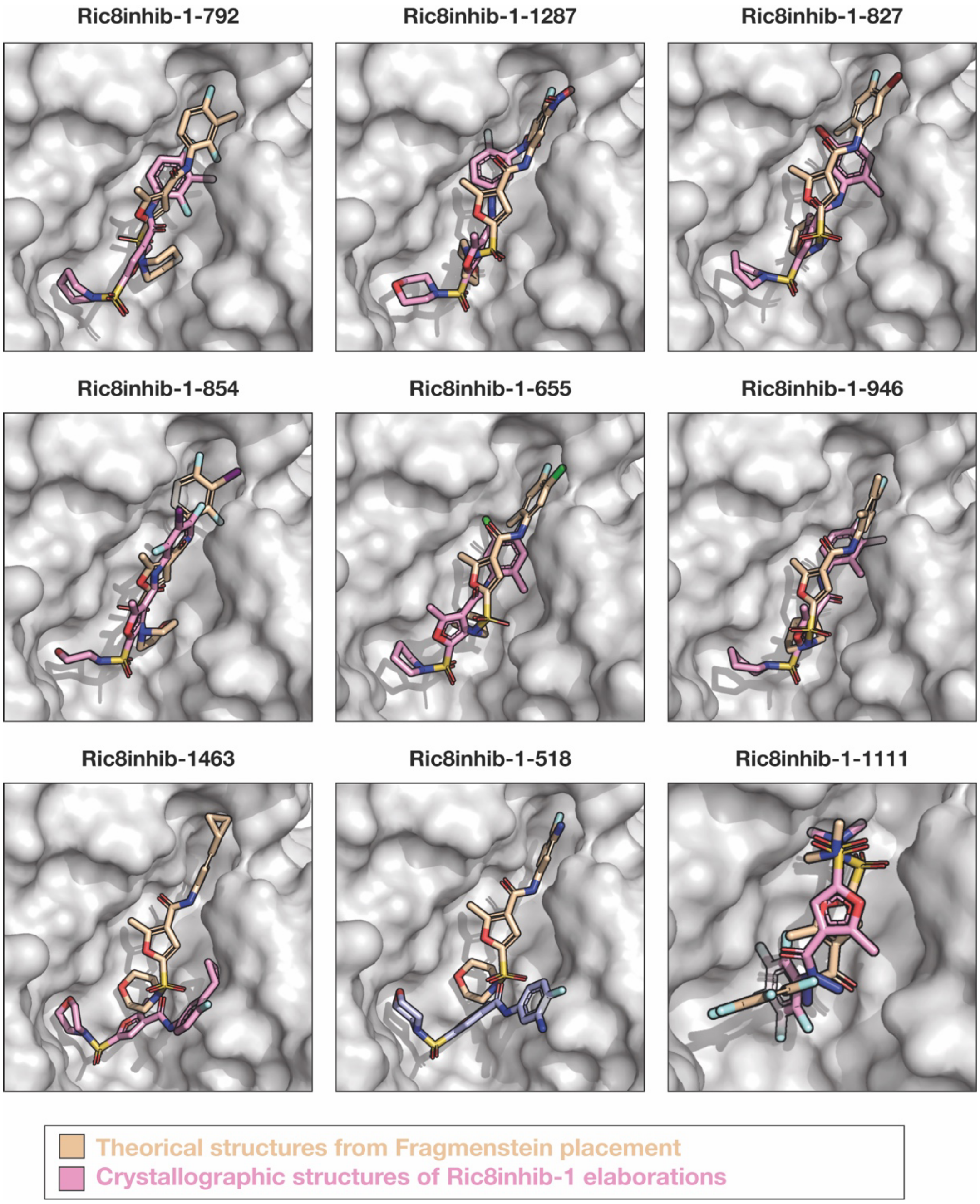


**Figure S11. Superimposition of crystallographic structures of Ric8inhib-1 elaborations with poses predicted by Fragmenstein.** The proposed placements by Fragmenstein, based on fragment structures, are shown as light orange sticks. The crystallographic structures of Ric8inhib-1 elaborations are displayed as pink sticks.

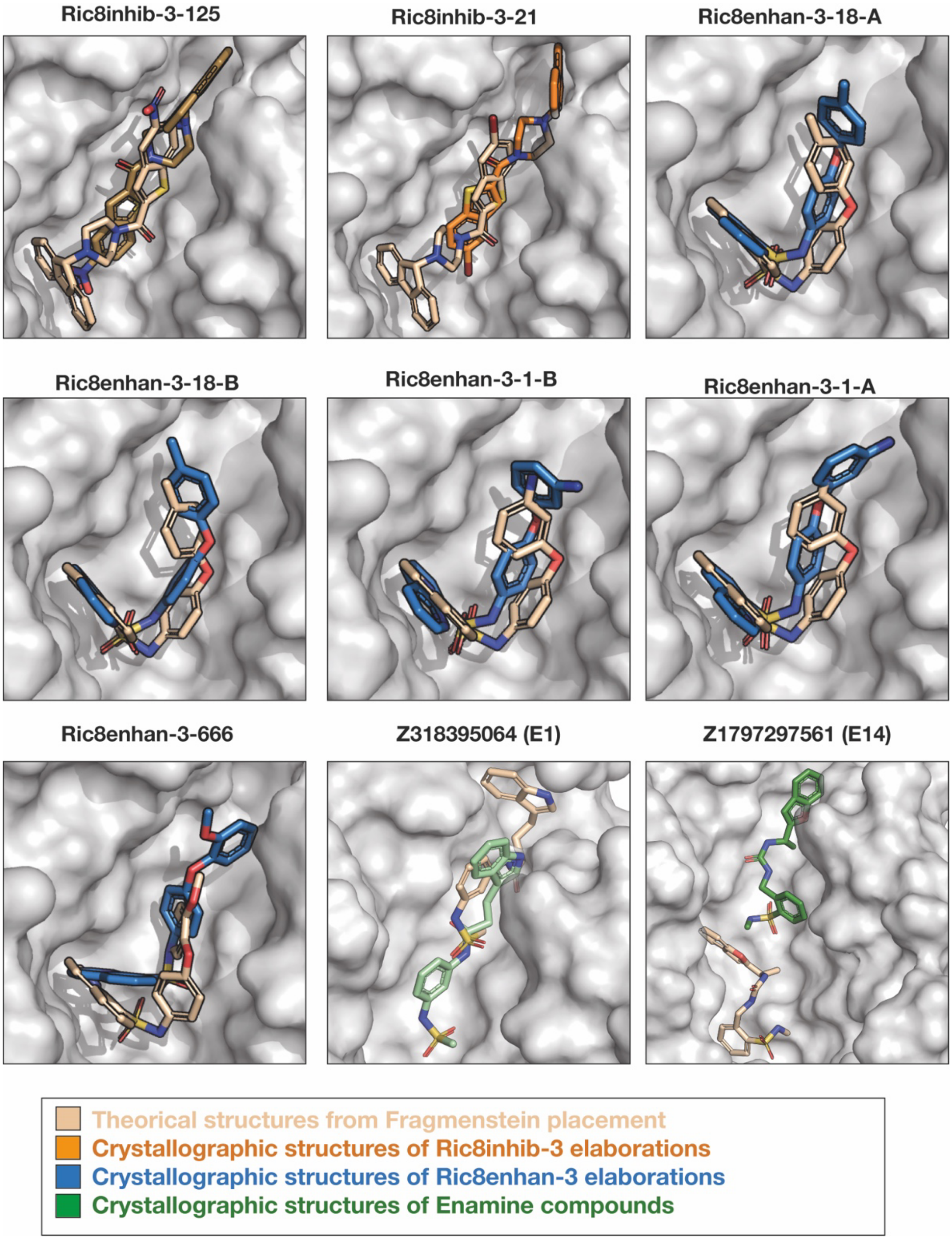


**Figure S12. Superimposition of crystallographic structures of various elaborations relative to placements predicted by Fragmenstein.** The placements proposed by Fragmenstein, based on fragment structures, are shown as light orange sticks. The crystallographic structures of Ric8inhib-3 elaborations are displayed as orange sticks, Ric8enhan-3 elaborations as blue sticks, and catalog compounds as green sticks.

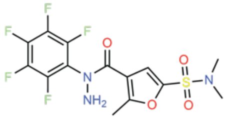

**Ric8A-inhib-1-1111**
NCS1-x1888
Ligand A – Chain A

Fragments: x0119-2B
x0311-1A
RMSD: 3.42 Å

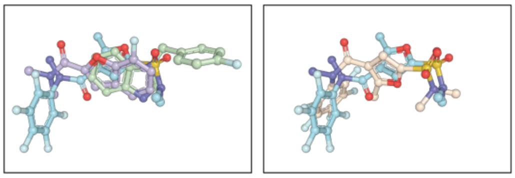

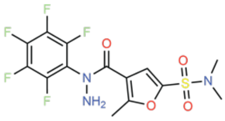

**Ric8A-inhib-1-1111**
NCS1-x1888
Ligand B – Chain A

Fragments: x0119-2B
x0311-1A
RMSD: 8.29 Å

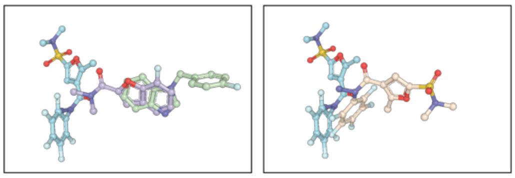

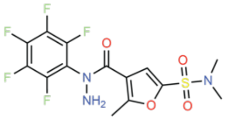

**Ric8A-inhib-1-1111**
NCS1-x1888
Ligand A – Chain B

Fragments: x0119-2B
x0311-1A
RMSD: 8.23 Å

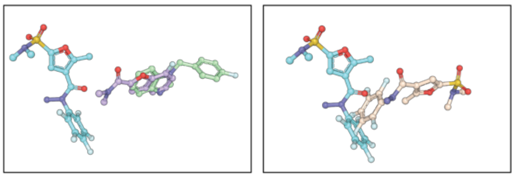

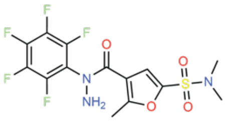

**Ric8A-inhib-1-1111**
NCS1-x1888
Ligand B – Chain B

Fragments: x0119-2B
x0311-1A
RMSD: 3.80 Å

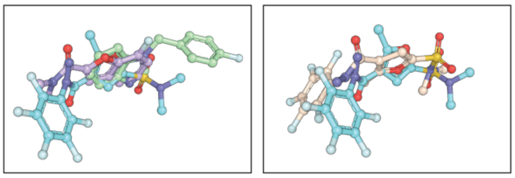

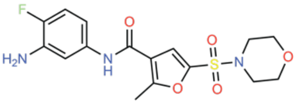

**Ric8A-inhib-1-518**
NCS1-x2005
Chain B

Fragments: x0119-2B
x0311-1A
RMSD: 8.38 Å

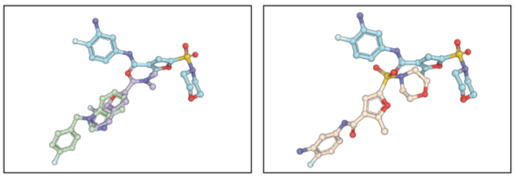

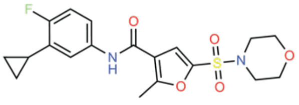

**Ric8A-inhib-1-1463**
NCS1-x1740
Chain B

Fragments: x0119-2B
x0311-1A
RMSD: 8.76 Å

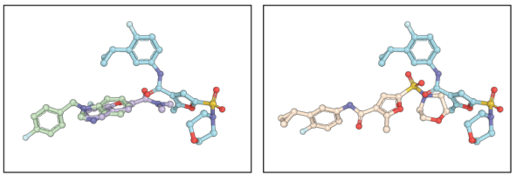

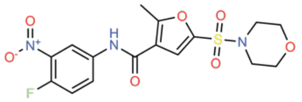

**Ric8A-inhib-1-1287**
NCS1-x1696
Chain A

Fragments: x0119-2B x0311-1A RMSD: 4.23 Å

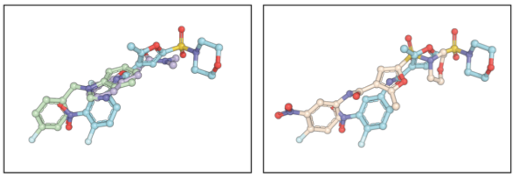

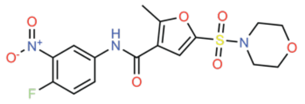

**Ric8A-inhib-1-1287**
NCS1-x1696
Chain B

Fragments: x0119-2B x0311-1A RMSD: 4.34 Å

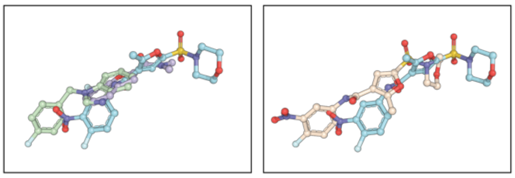

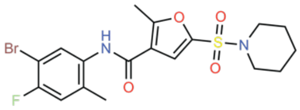

**Ric8A-inhib-1-827**
NCS1-x1943
Chain A

Fragments: x0119-2B x0311-1A RMSD: 4.60 Å

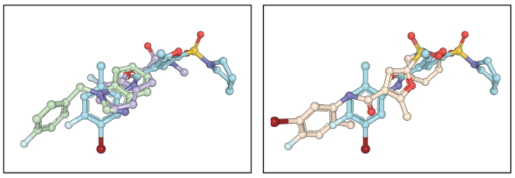

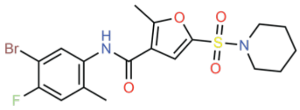

**Ric8A-inhib-1-827**
NCS1-x1943
Chain B

Fragments: x0119-2B x0311-1A RMSD: 4.47 Å

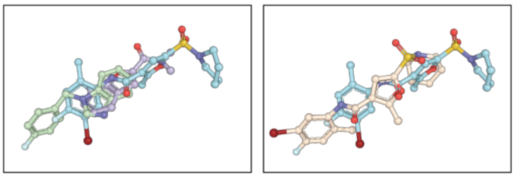

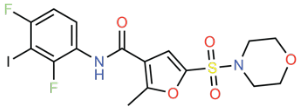

**Ric8A-inhib-1-854**
NCS1-x1726
Chain B

Fragments: x0119-2B x0311-1A RMSD: 4.47 Å

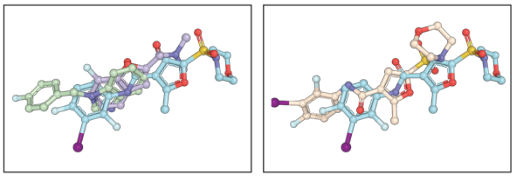

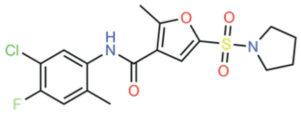

**Ric8A-inhib-1-655**
NCS1-x1894
Chain A

Fragments: x0119-2B x0311-1A RMSD: 5.19 Å

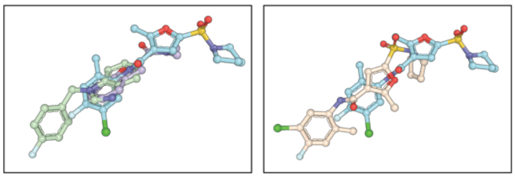

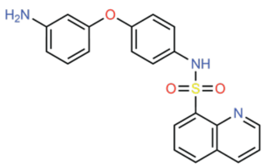


**Ric8A-enhan-3-1**
NCS1-x0975
Chain A

Fragments: x0119-2B x0311-1A RMSD: 2.76 Å

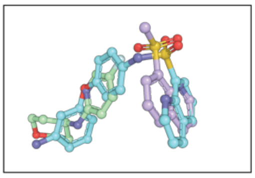
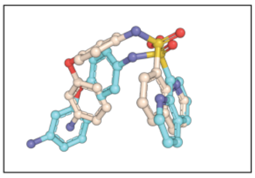

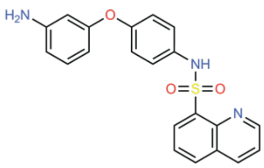


**Ric8A-enhan-3-1**
NCS1-x0975
Chain B

Fragments: x0119-2B x0311-1A RMSD: 2.93 Å

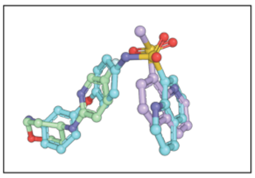
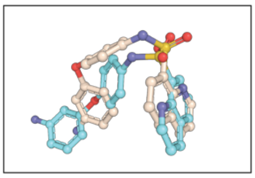

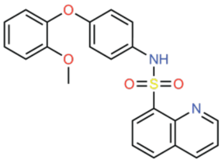


**Ric8A-enhan-3-666**
NCS1-x0981
Chain A

Fragments: x0071-0B x0119-2B RMSD: 3.26 Å

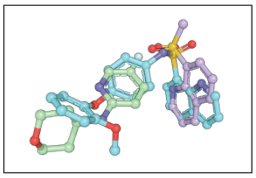
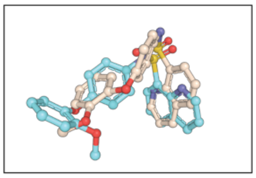

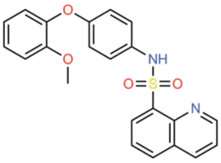


**Ric8A-enhan-3-666**
NCS1-x0981
Chain B

Fragments: x0071-0B x0119-2B RMSD: 4.22 Å

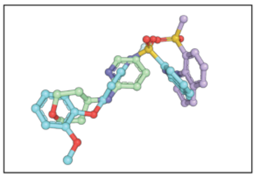
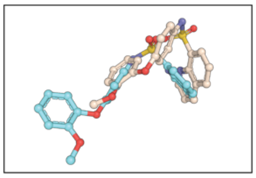

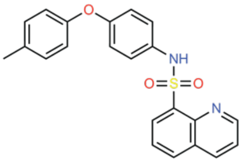


**Ric8A-enhan-3-18**
NCS1-x0974
Chain B

Fragments: x0125-0A x0580-0A RMSD: 2.52 Å

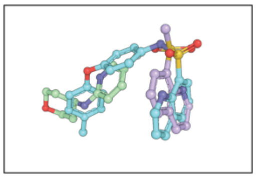
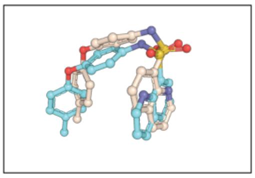

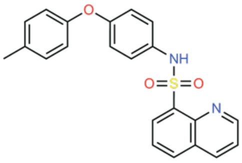


**Ric8A-enhan-3-18**
NCS1-x0974
Chain A

Fragments: x0125-0A x0580-0A RMSD: 2.90 Å

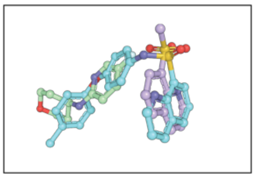
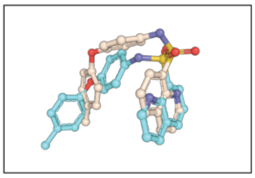

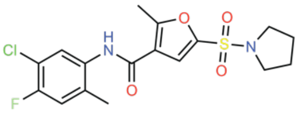

**Ric8A-inhib-1-655**
NCS1-x1894
Chain B

Fragments: x0119-2B x0311-1A RMSD: 4.97 Å

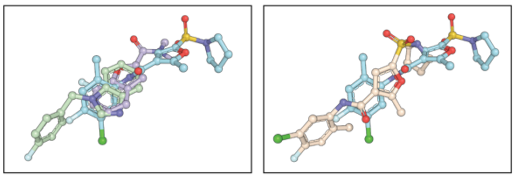

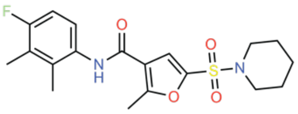

**Ric8A-inhib-1-946**
NCS1-x1942
Chain A

Fragments: x0119-2B x0311-1A RMSD: 3.82 Å

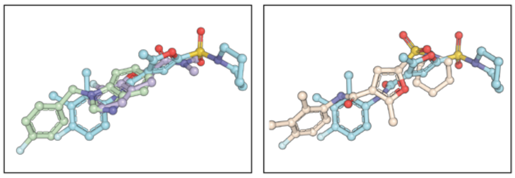

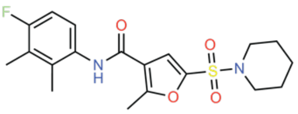

**Ric8A-inhib-1-946**
NCS1-x1942
Chain B

Fragments: x0119-2B x0311-1A RMSD: 4.16 Å

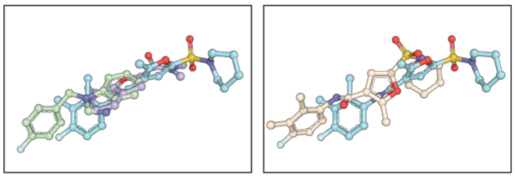

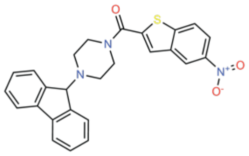

**Ric8A-inhib-3-125**
NCS1-x1698
Chain A

Fragments: x0071-0B x0119-2B RMSD: 10.94 Å

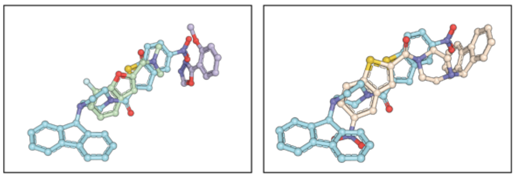

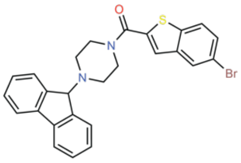

**Ric8A-inhib-3-21**
NCS1-x1699
Chain B

Fragments: x0119-2B x0071-0B RMSD: 12.16 Å

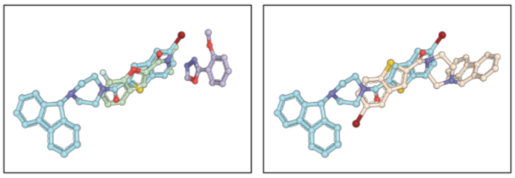

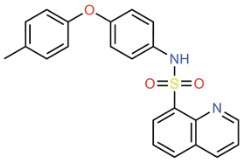

**Ric8A-enhan-3-18**
NCS1-x0974
Chain A

Fragments: x0125-0A x0580-0A RMSD: 2.90 Å

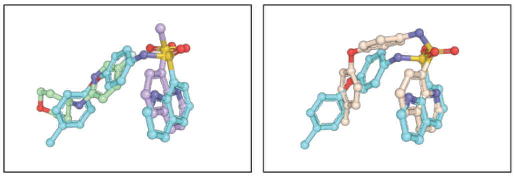

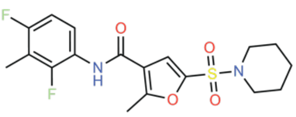

**Ric8A-inhib-1-792**
NCS1-x1238
Chain A

Fragments: x0119-2B, x0311-1A    RMSD: 5.49 Å

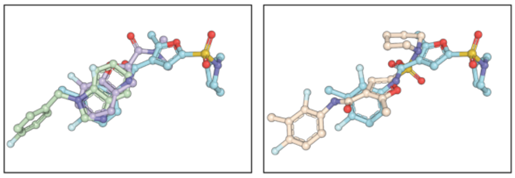

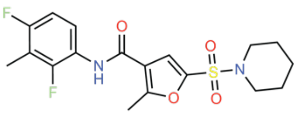

**Ric8A-inhib-1-792**
NCS1-x1238
Chain B

Fragments: x0119-2B, x0311-1A    RMSD: 4.85 Å

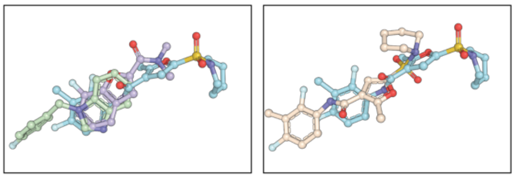

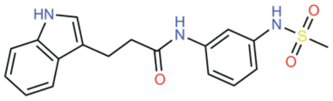

**E1 (Z318395064)**
NCS1-x1748
Chain A

Fragments: x0110-2B, x0252-2B    RMSD: 15.57 Å

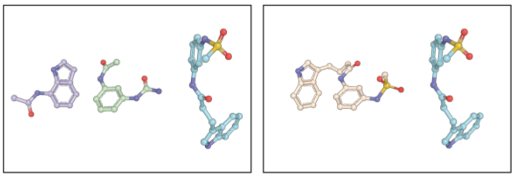

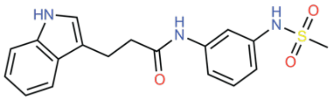

**E1 (Z318395064)**
NCS1-x1748
Chain B

Fragments: x0110-2B, x0252-2B    RMSD: 5.87 Å

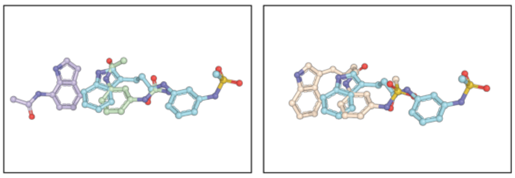

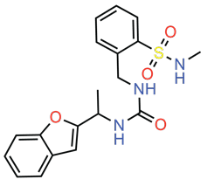

**E14 (Z1797297561)**
NCS1-x1777
Chain A

Fragments: x0119-0B, x0469-0B    RMSD: 12.20 Å

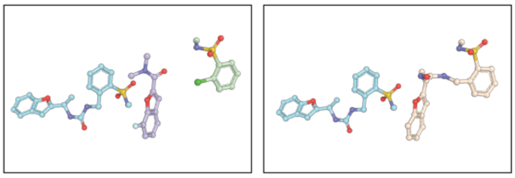

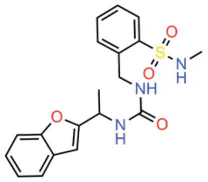

**E14 (Z1797297561)**
NCS1-x1777
Chain B

Fragments: x0119-0B, x0469-0B    RMSD: 12.55 Å

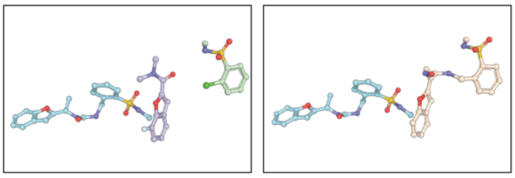

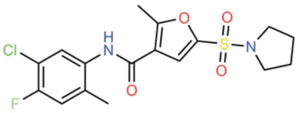

**Ric8A-inhib-1-655**
NCS1-x1894
Chain B

Fragments: x0119-2B x0311-1A RMSD: 4.97 Å

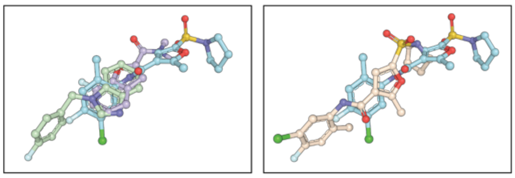

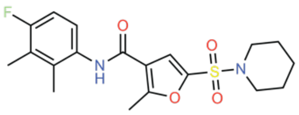

**Ric8A-inhib-1-946**
NCS1-x1942
Chain A

Fragments: x0119-2B x0311-1A RMSD: 3.82 Å

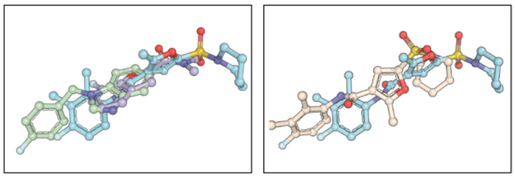

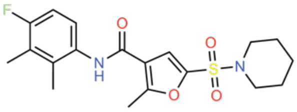

**Ric8A-inhib-1-946**
NCS1-x1942
Chain B

Fragments: x0119-2B x0311-1A RMSD: 4.16 Å

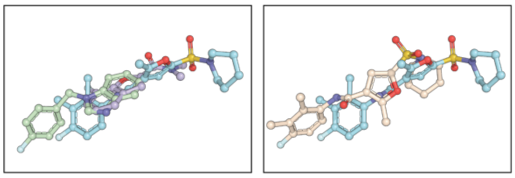

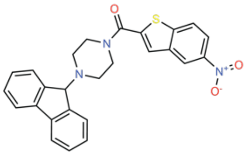

**Ric8A-inhib-3-125**
NCS1-x1698
Chain A

Fragments: x0071-0B x0119-2B RMSD: 10.94 Å

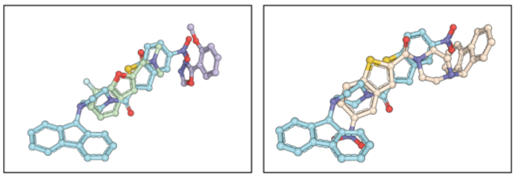

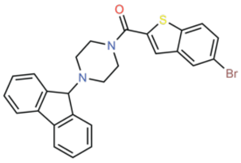

**Ric8A-inhib-3-21**
NCS1-x1699
Chain B

Fragments: x0119-2B x0071-0B RMSD: 12.16 Å

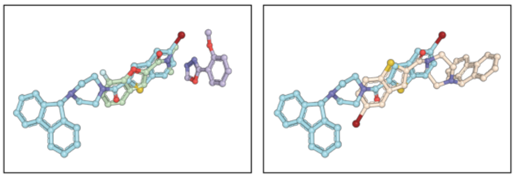

**Figure S13. Superimposition of crystallographic structures of various elaborations relative to crystallographic structures of their inspiration fragments.** For each compound resolved, the 2D structure, name, Fragalysis name, and chain are labeled. The inspiration fragments for the design are listed and the resolved structures of the fragments (purple and green) are overlaid with the resolved follow-up compound (blue). The RMSD to the predicted pose (cream) is listed and is shown in relation to the resolved structure (blue).

### Supplementary tables

**Table S1: Selected pure catalog compounds.** (*) Indicates the Site (S) where the inspiration fragments were found in the fragment screening. (**) Hydrogen bonds, π-π interactions, and hydrophobic interactions were considered.

| | | ***Fragmenstein*** | | | **Predicted key interactions (**)** | | | | | | | |
|---|---|---|---|---|---|---|---|---|---|---|---|---|
| **Pure compounds** | **Fragments (*)** | **ΔG (kcal/mol)** | **RMSD (Å)** | **Tanimoto** | **W30 (**)** | **D37** | **Y52** | **T92** | **K100** | **W103 (**)** | **Y108** | **R148** |
| Z318395064 (E1) | Z1255459547 (S1) + Z198195770 (S2) | -10.1 | 0.78 | 0.36 | X | X | X | X | | | | |
| Z2005021751 (E2) | Z1255459547 (S1) + Z133716556 (S2) | -10.9 | 0.83 | 0.42 | | | X | X | | X | | |
| Z451568306 (E3) | Z1203730757 (S4) + Z54226006 (S7) | -9.7 | 0.44 | 0.32 | | | | | | | | X |
| Z1097155203 (E4) | Z239127534 (S1) + Z26333448 (S2) | -8.0 | 0.50 | 0.45 | X | X | | X | | | | |
| Z57379493 (E5) | Z55290386 (S3) + Z1269220427 (S4) | -12.0 | 0.88 | 0.48 | | | X | X | | X | | |
| Z1260603230 (E6) | Z552903867 (S3) + Z126922042 (S4) | -12.0 | 0.88 | 0.48 | | | X | X | | X | | |
| Z1104573517 (E7) | Z1255459547 (S1) + Z198195770 (S2) | -12.0 | 0.79 | 0.33 | X | X | X | | | | | |
| Z1556583850 (E8) | Z300081412 (S5) + Z1272480091 (S6) | -9.5 | 0.68 | 0.38 | | | X | | | X | X | |
| Z952525226 (E9) | Z29692148 (S1) + Z26333448(S2) | -7.2 | 0.64 | 0.58 | X | X | | | | | | |
| Z1396906847 (E10) | Z26333448 (S2) + Z1265813904 (S3) | -9.6 | 0.99 | 0.28 | | | X | X | | | | |
| Z1367764037 (E11) | POB0063 (S5) + Z1272480091 (S6) | -11.4 | 0.99 | 0.34 | | | | X | | | | |
| Z1870241397 (E12) | Z166605460 (S4) + Z1272480091(S6) | -11.0 | 0.99 | 0.24 | | | | | X | | | |
| Z2234189231 (E13) | Z31723394 (S2) + Z55290386 (S3) | -9.7 | 0.92 | 0.23 | X | X | X | X | | X | | |
| Z2234189231 (E14) | Z1265813904 (S4) + Z45527714 (S7) | -12.8 | 0.99 | 0.38 | | | X | X | | | | |
| Z1798494323 (E15) | Z1255459547 (S4) + Z300081412 (S5) | -11.1 | 0.85 | 0.35 | | | X | | | | | |
| Z3401857786 (E16) | Z1255459547 (S1) + Z26333448 (S2) | -7.5 | 0.96 | 0.62 | X | X | | X | | | | |
| Z7823261935 (E17) | Z1198177230 (S4) + Z32327641 (S5) | -9.3 | 0.61 | 0.33 | | | | | | X | | X |

**Table S2: Comparison of affinity results obtained via GCI and Trp fluorescence emission techniques.**

| Pure compound | $K_d$, (μM) GCI | $K_d$, (μM) Trp fluorescence emission |
|---|---|---|
| Z1260603230 (E6) | 63.71 | 27.5 ± 5.1 |
| Z1396906847 (E10) | 97.31 | 398.5 ± 98.6 |
| Z1870241397 (E12) | 103.74 | 34.5 ± 15.0 |
| Z2234189231 (E14) | 102.82 | 77.62 ± 0.02 |
| Z3401857786 (E16) | 110.23 | 98.5 ± 5.6 |
| Z7823261935 (E17) | 141.45 | 263.4 ± 44.8 |

**Table S3: Summary of the different steps during GCI for NCS-1 capture on the NTA surface.**

| Step | Solution | Concentration | Time (s) | Flux (μl/min) |
|---|---|---|---|---|
| Activation | $NiCl_2$ | 500 μM | 60 | 30 |
| Wash | EDTA | 3 mM | 60 | 30 |
| Capture | His-NCS-1 | 15 μg/ml | 60 | 10 |
| Sample | Pure compounds | 100 μM | 25 | 100 |
| Regeneration | EDTA | 350 mM | 60 | 30 |
